# Automated 3D Landmarking of the Skull: A Novel Approach for Craniofacial Analysis

**DOI:** 10.1101/2024.02.09.579642

**Authors:** Franziska Wilke, Harold Matthews, Noah Herrick, Nichole Dopkins, Peter Claes, Susan Walsh

## Abstract

Automatic dense 3D surface registration is a powerful technique for comprehensive 3D shape analysis that has found a successful application in human craniofacial morphology research, particularly within the mandibular and cranial vault regions. However, a notable gap exists when exploring the frontal aspect of the human skull, largely due to the intricate and unique nature of its cranial anatomy. To better examine this region, this study introduces a simplified single-surface craniofacial bone mask comprising 9,999 quasi-landmarks, which can aid in the classification and quantification of variation over human facial bone surfaces.

Automatic craniofacial bone phenotyping was conducted on a dataset of 31 skull scans obtained through cone-beam computed tomography (CBCT) imaging. The MeshMonk framework facilitated the non-rigid alignment of the constructed craniofacial bone mask with each individual target mesh. To gauge the accuracy and reliability of this automated process, 20 anatomical facial landmarks were manually placed three times by three independent observers on the same set of images. Intra- and inter-observer error assessments were performed using root mean square (RMS) distances, revealing consistently low scores.

Subsequently, the corresponding automatic landmarks were computed and juxtaposed with the manually placed landmarks. The average Euclidean distance between these two landmark sets was 1.5mm, while centroid sizes exhibited noteworthy similarity. Intraclass coefficients (ICC) demonstrated a high level of concordance (>0.988), and automatic landmarking showing significantly lower errors and variation.

These results underscore the utility of this newly developed single-surface craniofacial bone mask, in conjunction with the MeshMonk framework, as a highly accurate and reliable method for automated phenotyping of the facial region of human skulls from CBCT and CT imagery. This craniofacial template bone mask expansion of the MeshMonk toolbox not only enhances our capacity to study craniofacial bone variation but also holds significant potential for shedding light on the genetic, developmental, and evolutionary underpinnings of the overall human craniofacial structure.

## INTRODUCTION

The field of phenomics – understanding the qualitative and quantitative traits that characterize a phenotype-is a fast-developing field [1]. Over the past two decades, numerous publications have not only unveiled genetic variants associated with phenotypes, but also made significant advancements in phenotyping methodologies [2, 3]. Moreover, the emergence of new technologies has enabled us to capture high quality 3D scans, encompassing both hard and soft tissue structures [1, 4, 5]. Although there have been significant strides made in understanding facial soft tissue variation, with technical advances implemented for genome wide association studies (GWAS) on facial shape [3, 6], the underlying craniofacial structure remains largely unexplored. This is in part due to the intricate nature of the entire skull shape and challenges in acquiring large numbers of 3D scans. Nevertheless, understanding human craniofacial structure is pivotal due to its substantial contribution to our facial appearance, particularly owing to its relative independence from biological factors such as weight and reduced susceptibility to agerelated changes after reaching adulthood [7, 8]. Hence, a comprehensive exploration of skull morphology is essential for gaining a holistic understanding of the genetic determinants governing human facial shape. Although a recent GWAS was performed on the cranial vault [9], a more comprehensive study of the viscerocranium (craniofacial bone structure) is imperative to tie in with facial soft tissue research that has been so successful in recent years.

Typically, studies describing the shape of the human skull have predominantly been within the field of Anthropology. In this context, the shape of the skull has been used to categorize an individual’s sex and ancestral origins [10, 11]. While sex often relies on visual indicators, the assessment of ancestry is more complex. Computational tools such as FORDISC [12] utilize skull measurements to estimate ancestry but cannot account for admixture and smaller subpopulations. Furthermore, estimations of soft-tissue thickness have been employed for facial reconstruction from a skull [13, 14] and skull shape analyses provide insights on primates to *homo sapiens* evolutionary processes [15, 16]. In the medical realm, skull shape is often used to describe specific pathologies or act as a non-syndromic reference [17, 18]. More recently, the dental and plastic surgery fields have also capitalized on skull shape analyses to aid in reconstructive surgical procedures [19, 20]. However, many of these previous approaches were constrained by the reliance on manual cranial landmarks (usually less than 50) and the use of physical skulls or radiographs as data sources [21, 22]. These conventional approaches bring inherent challenges. Firstly, the process is time-consuming, requires trained observers, and is prone to intra- and inter-observer error, thereby complicating standardization [23-25]. Although shape analysis can be performed by considering the overall configuration of a few landmarks, such an approach trivializes the complete complexity of cranial facial shape [26, 27]. Although, algorithms were developed in the early 2000s that have allowed automatic 3D dense phenotyping [28], they have vastly improved since then [26, 27, 29].

A more recent framework for automatic 3D dense phenotyping, “MeshMonk”, was introduced by White et al. [27] in 2019. MeshMonk provides a facial soft tissue mesh comprising approximately 7160 points, accompanied by algorithms to facilitate the alignment of this mask to 3D facial scans. This framework has provided a straightforward, standardized, and validated method to describe the phenotypic variation found in facial shape using large datasets by simplifying automated landmarking. While its application has led to multiple publications exploring the genetic architecture of the human face [2, 3, 6], it is crucial to note however, that the framework does not include a complete hard-tissue component. Global registration masks using Meshmonk have been developed and utilized for specific segments of the skull, including the lower jaw [30] and the cranial vault [9], however, assessment of the facial bones within the craniofacial complex has not been explored. The frontal aspect of the skulls facial skeleton poses significant challenges due to its intricate structure, with a separate lower jaw, as well as orbital and nasal cavities. While the performance of a full skull mask application using the meshmonk framework has already been published [31], the mask itself is not freely available for download, it also utilizes approximately 155,000 vertices making it computationally intensive, in addition to the full cranium being a specific requirement that is often not present in CBCT scans. This prevents its use on dental CBCT scans which are one of the most common types of facial scans performed. To perform large-scale GWAS that explore hard structures of the human face, it requires a large number of bone scans to be processed accurately and efficiently.

Fortunately, advancements in medical technology have increased the availability of these scans via Magnetic Resonance Imaging (MRI) and Computer Tomography (CT). The use of Cone Beam CT (CBCT) has also emerged as a prominent imaging technique in the dental field. CBCT scans have the advantage of lower radiation dosages than conventional CT scans and reduced costs, rendering them viable for research purposes. To facilitate and cover the broad range of bone scans available, a suitable landmarking approach must be devised to simplify the intricate aspects of skull morphology without sacrificing critical information, therefore ensuring a more comprehensive and powerful analysis.

For this research, by focusing solely on facial bones within the frontal region of the skull, we provide a simplified mesh which reduces computational demands that is compatible with the MeshMonk framework. The frontal region of the skull encompassing the facial bones is captured and reconstructed as a 3D replicate. A template or “mask” consisting of thousands of points (n=9999), or “quasi-landmarks” are aligned and non-rigidly mapped onto the target following the targets geometry. These quasi-landmarks replace manually placed positions and are in anatomical correspondence across all individuals, allowing a more comprehensive shape evaluation. Validation, reliability, and accuracy assessments of the quasi-landmark placement is accomplished by comparing the automatically placed landmarks with manually positioned landmarks. The present research constitutes an important extension to the MeshMonk framework, enabling its application to skull scans, both CT and CBCT, thereby empowering researchers to delve more easily into the analysis of craniofacial bone structure. This augmentation broadens the potential scope of investigations in phenomics research and facilitates a comprehensive exploration of the genetic determinants underlying the human skull, in particular craniofacial bone morphology.

## MATERIALS & METHODS

### PARTICIPANT RECRUITMENT & STUDY SAMPLE

Participants for this research were collected at Indiana University Indianapolis (IUI). The study underwent ethical review and received approval from the institutional review board (IU IRB 1801992304). Prior to participating, individuals provided informed consent, which included disclosure of potential radiation exposure associated with Cone Beam Computed Tomography (CBCT) imaging. To ensure anonymity, each participant was assigned a unique identification number, and all collected data were securely stored on a server accessible only to those with pre-existing ethical authorization. The study exclusively enrolled individuals aged 18 and above, excluding those with a history of significant facial trauma, individuals with incomplete data, or scans that lacked complete orbital information. In total, the dataset comprised 31 skulls.

CBCT imaging procedures were conducted at the IU School of Dentistry within the Orthodontics and Oral Facial Genetics Department, utilizing a Carestream 9300 machine manufactured by Carestream Health, Inc. (NY). All full-face scans adhered to specific parameters, including a field of view (FOV) of 17 cm x 13.5 cm (this encompasses all facial bones, whilst excluding most of the frontal bone/ forehead), an X-ray tube current of 15 mA, an X-ray tube voltage of 90 kV, and a scan duration of 28 seconds. The scans themselves were administered by a qualified and licensed professional.

### DICOM EXTRACTION AND DATA CLEANING

The DICOM images obtained from the CBCT were processed in the free software 3D Slicer [32]. A threshold of between 400-600 and maximum Hounsfield units was set. The resulting mesh was filtered for the largest island to remove pieces of the spine and loose internal structures. In addition, minor holes were closed using the closing smoothing with a kernel size of 2.0mm. The skull meshes were then imported into Blender [33] where a half-cylindrical mesh was placed around the skull and the shrink-wrap modifier applied (Supplementary Figure S1). In addition, the subdivision surface modifier was applied to increase the resolution. This process was repeated a further 5x with two decimate modifiers (un-subdivide setting) in-between to prevent the resulting mesh file from being too large. The output was a high polygon count mesh with an uneven vertex distribution. To counteract this, the meshes were reduced to 30,000 triangles in Meshmixer [34] then evenly re-meshed, resulting in regularly spaced vertices, and a similar number of faces between meshes.

### PHENOTYPING

The most even and complete skull mesh was used as a preliminary mask and symmetrized in Blender (one half delete, the center vertices moved to X=0 and the mesh mirrored). A subset of 20 skulls were masked with this preliminary mask using the Meshmonk framework [27] in Matlab [35]. This toolbox uses a 3-step process to non-rigidly align the mask to each target shape: 1) initialization is performed by placing eleven manual landmarks (Supplementary Figure S2) (custom script within MeVisLab (available: http://www.mevislab.de/)) on both the mask and target shapes which are utilized to estimate the rigid registration, 2) rigid registration is optimized via iterative closest point registration, 3) non-rigid registration is performed to adapt the shape of the mask to the shape of the target mesh. Thus, resulting in all 20 skulls consisting of 9,999 quasi-landmarks in corresponding anatomical locations. After Procrustes Superimposition for alignment and scaling, the skulls were averaged resulting in the final average mask. This mask was consequently symmetrized in shape by averaging the original and its reflection.

The full set of 31 skulls were then registered via Meshmonk using the final average mask and eleven initial landmarks. This process was repeated three times with three different sets of initial landmarks to test the reliability of the automatic landmarking using the newly developed bone mesh. The process from DICOM data to masked skull took approximately 30 minutes per skull, however this time depends predominantly on the speed of the CPU.

### VALIDATION LANDMARKS

To assess the agreement between manually and automatically placed landmarks, the 31 skulls were landmarked manually with 20 landmarks by three observers (Figure 1). Observer 1 was well versed in this procedure due to their anthropological background; the others were untrained. Landmarks were chosen which evenly covered the skull and represented the areas consistently captured by CBCT with clear definitions taken from literature [31, 37]. These “gold-standard” landmarks were placed on the skulls after shrink-wrapping and remeshing as described previously for a more accurate comparison. Each observer landmarked all 31 skulls three times, with at least 24 hours between sessions.

**Figure 1:**
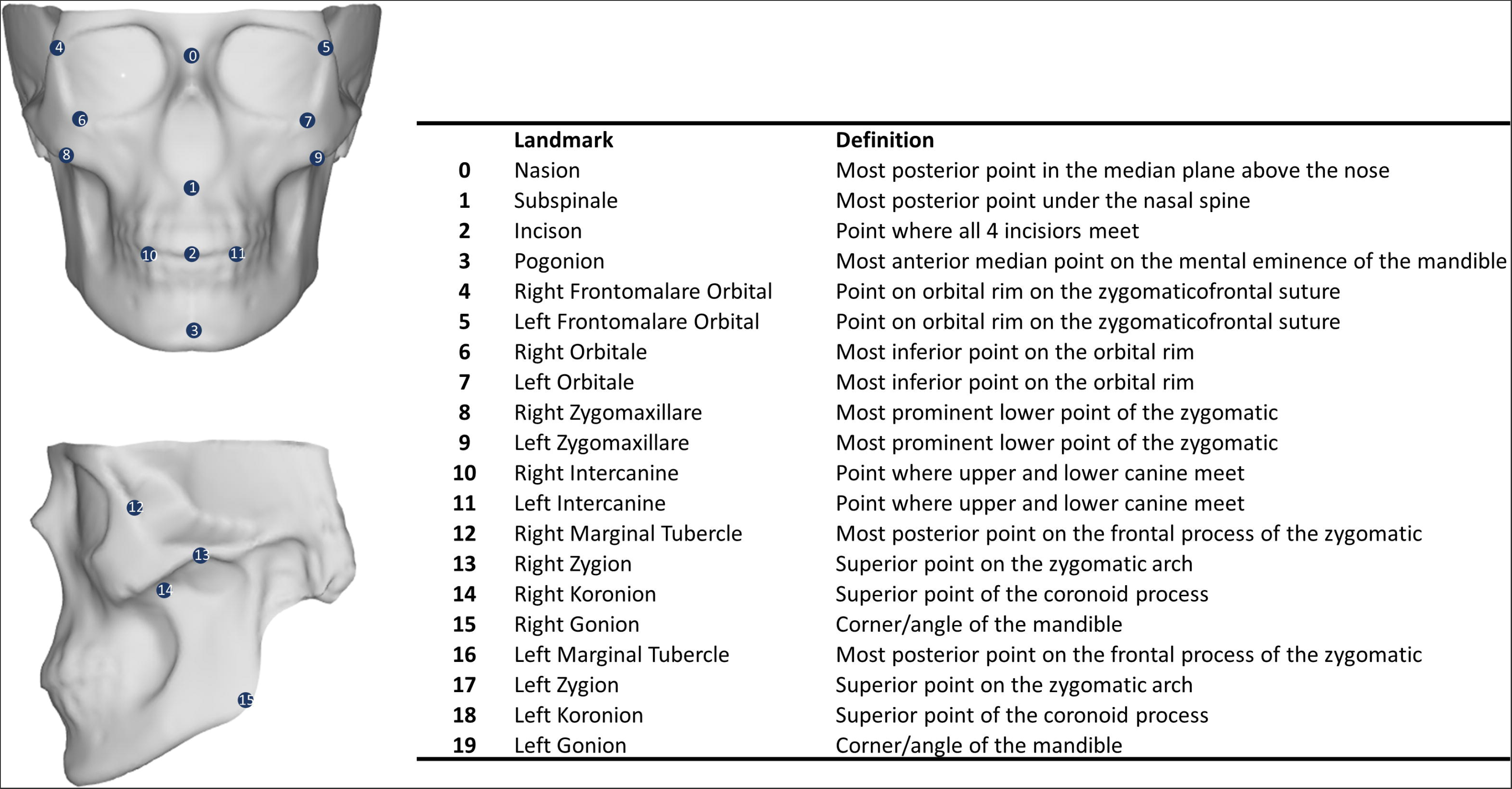
Overview of the manually placed landmarks. Landmark definitions were taken from [31, 37].

To automatically place landmarks which coincide with the manual landmarks, a leave-one-out approach was used. One skull was determined to be the target skull, while the remaining 30 were the training dataset. Manually placed landmarks (averages per observer over the 3 landmarking rounds) were transferred to the masked skulls by translating them to barycentric coordinates. Their location on the masked skull was calculated via a weighted sum (Barycentric coordinates) of the three closest quasi-landmarks. These were averaged over the training set and then translated back to cartesian coordinates on the target skull. Due to the process of averaging, the resulting landmark on the target skull was not always on the surface. To circumvent this issue, the landmark was projected to the closest point on the surface of the target skull. This placement was repeated using each observers’ manual landmarks individually, and an average of all observers’ landmarks.

An overview of the pre-assessment process can be found in Figure 2.

**Figure 2:**
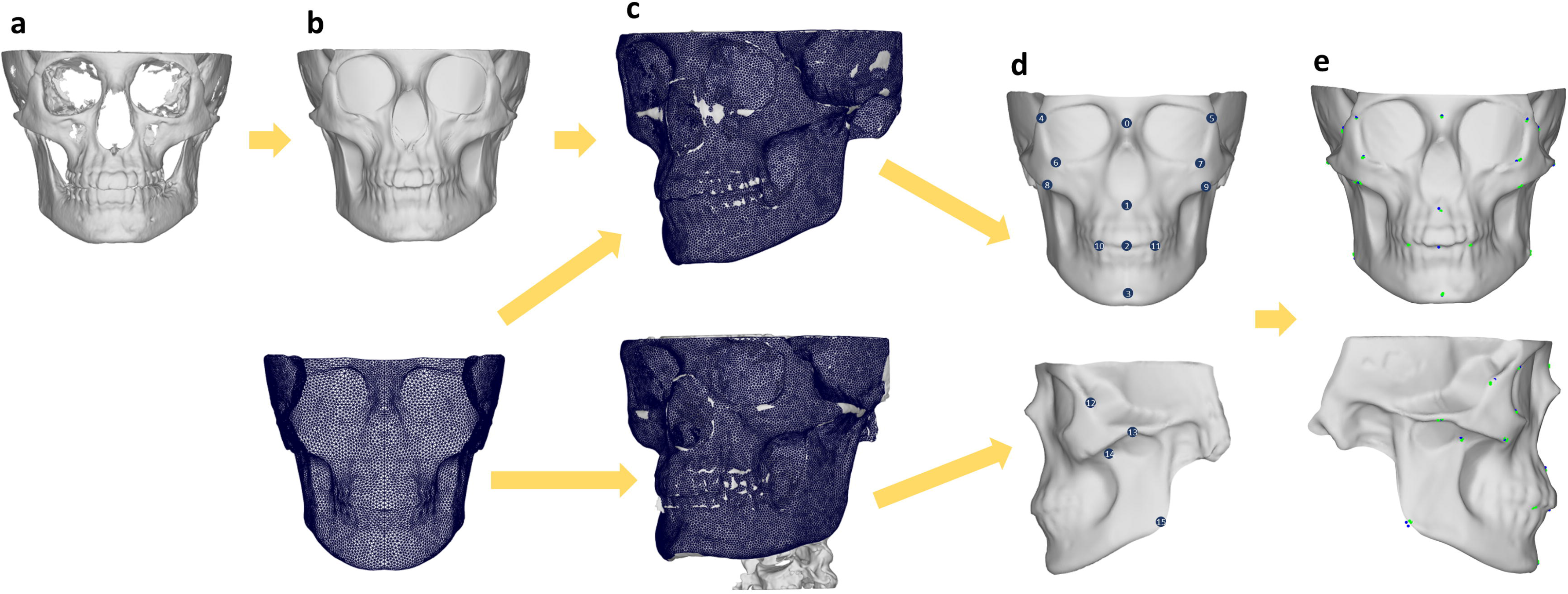
Overview of pre-assessment steps. A: Skull mesh is exported from 3D slicer, B: Skull mesh is shrink wrapped in Blender, C: craniofacial bone mask is applied to all skulls using Meshmonk, D: 20 Landmarks are placed manually by 3 observers, E: Automatic Landmarks are determined.

### RELIABILITY ASSESSMENT

While automatic landmarking is consistent within itself, the MeshMonk toolbox requires the placement of initial landmarks for registration. As these are placed manually, variation can be present. Reliability of this process was assessed by calculating the Root Mean Square (RMS) distance (root square of the mean of squared Euclidean distances) of the resulting masked quasi-landmarks to the centroid (mean point over all landmarks) over all three iterations. A smaller RMS shows less variation between quasi-landmark placement. RMS was calculated for both the 20 automatic landmarks, as well as the 9,999 quasi-landmarks on the craniofacial bone mask.

Intra-Observer reliability was calculated as the RMS between the three rounds of landmarking for each observer, the centroid size of the landmark configuration, as well as the standard deviation of the xyz coordinates separately. Inter-observer reliability was between the three observers over the average of their three landmarking rounds. The reliability of the automatic landmarking was analyzed using the three sets of automatic landmarks derived from the three observers’ manual landmarks.

To analyze if the automatic and manual landmarks were more or less variable, descriptive statistics (Mean, Standard Deviation, Minimum, Maximum) as well as an ANOVA on centroid size with Observer, Skull, and nested Observer/Iteration was performed. MANOVA were performed on the generalized Procrustes analysis (GPA) aligned landmarks for both the manual and automatic landmarks with Skull and Observer as factors (as well as nested Observer/Iteration for the manual landmarks) to see which of these explained variation within the landmarks. In addition, Levene’s Test was performed on the variance of the standard deviation over the xyz coordinates between the automatic and manual landmarks to analyze if the error variation was statistically different. Intraclass correlation coefficients (ICC) were calculated for intra and inter-observer centroid sizes (two-way consistency (inter-) and agreement (intra-).

All statistical analyses were performed either in Matlab, or in R using the packages Geomorph [38], irr [39], and SimplyAgree [40]. Plots were created using ggplot2 [41].

### ACCURACY ASSESSMENT

To calculate the accuracy of the automatic landmarks in relation to the ‘gold standard’ manual landmarks, multiple approaches were used. Initially, the Euclidean distance between the average manual and automatic landmarks were calculated to provide basic information as to which landmarks show the highest accuracy. Bland-Altman plots were used to visualize the agreement between centroid sizes, as well as individually for xyz coordinates. ICC statistics were used to compare landmark indications using both xyz coordinates and centroid sizes (two-way agreement). To determine if the method explained variation between the landmarks, an ANOVA was performed on centroid sizes with skull, observer, and method as predictors.

To determine which factors explained variance within the landmarks, multiple MANOVA tests were performed. The landmarks were GPA aligned and inputted into a MANOVA with skull, observer, and method as factors. A second MANOVA was performed on the principal component scores from a shape principal component analysis using the PCs which explained 95% of the variation between the landmarks.

Due to the shrink-wrapping process, gaps such as eye sockets are filled in and defined by quasi-landmarks. To calculate which of the landmarks are the best and most stable to define points on the physical skull we calculated the distance between the meshmonk skull and the original skull (before wrapping) along the normal vectors. Any landmarks with distances more than 10mm in more than half of the skulls were removed. The remaining skull was symmetrized so that the same landmarks were kept on either side.

### IMAGE APPLICATION

The craniofacial bone mask, instructions for CBCT export and shrink-wrapping, script for producing quality control images, as well as the IDs for the vertices that do not define true points on the skulls can be found in the Supplementary Material and on our website at https://walshlab.sitehost.iu.edu/pages/craniofacial.html.

We also provide visualization of a basic proof of application; our 31 masked skulls were used to analyze sexual dimorphism and perform a Principal Component Analysis (PCA) to show the variation attributed to sex and Principal Component (PC) 1. Each analysis was prefaced with a Partial Least Squares Regression (PLSR) to remove the effects of age, height, weight, ancestry, and sex (sex only to analyze PC1).

To analyze if this method could also be applied to CT images, we downloaded a CT image from the MUG500+ dataset [36]. These images were previously cleaned, and we ran a cleaned version through our masking pipeline.

## RESULTS

### RELIABILITY

Due to the fact that our “gold standard” landmarks were placed manually, a large emphasis must be made on the intra- and inter-observer error. Table 1 shows the average RMS distance over the 20 landmarks for each observer, inter-observer, and for the automated landmarking. RMS per landmark can be found in Supplementary Table 1 & 2. RMS distances were also calculated over the three meshmonk iterations for all 9,999 quasi-landmarks (Supplementary Figure 3) which show that the outer rim, especially over corners (mandibular angle, nasal spine), show the most variation in the automatic landmarking. However, the error at the gonion for automatic landmarking (0.33mm) is lower than that of manual landmarking (0.65mm), and smaller than the lowest error (Incisors = 0.41mm) for the trained observer. The majority of the central face has an error of under 0.1mm. The largest manual error is seen in the Zygomaxillare (2.22mm) for both inter- and intra-observer errors. On average, automatic landmarking was more than 5x more reliable than a trained observer. Variation in landmarking errors can be found in Supplementary Figure 4 & 5. In addition, the intra-observer standard deviation of the landmarking in the x, y, and z directions was calculated per landmark (Supplementary Table 3). The average standard deviation over all axes for the trained observer was 0.415mm with a range of 0.258-0.716mm. Observer 1 showed consistently smaller landmarking errors than the two untrained observers (Supplementary Figure S6). Intra-observer ICC was O1=0.998 (95% CI: 0.997 < ICC < 0.999), O2=0.988 (95% CI: 0.977 < ICC < 0.994) and O3=0.994 (95% CI: 0.987 < ICC < 0.997). Inter-observer ICC was 0.998 (95% CI: 0.997 < ICC < 0.999) showing high concordance in landmarking.

**Table 1:** RMS distances (in mm) of repeated landmarking averaged over the 20 landmarks. Automatic landmarking is the average over the three rounds of Meshmonk.

|  | Mean | Std | Min | Max |
| --- | --- | --- | --- | --- |
| **Automated** | 0.119 | 0.086 | 0.052 | 0.347 |
| **Inter-Observer** | 1.442 | 0.462 | 0.738 | 2.393 |
| **Intra-Observer 1** | 0.662 | 0.180 | 0.410 | 1.106 |
| **Intra-Observer 2** | 1.109 | 0.372 | 0.597 | 2.056 |
| **Intra-Observer 3** | 0.943 | 0.267 | 0.627 | 1.494 |

**Table 2:**
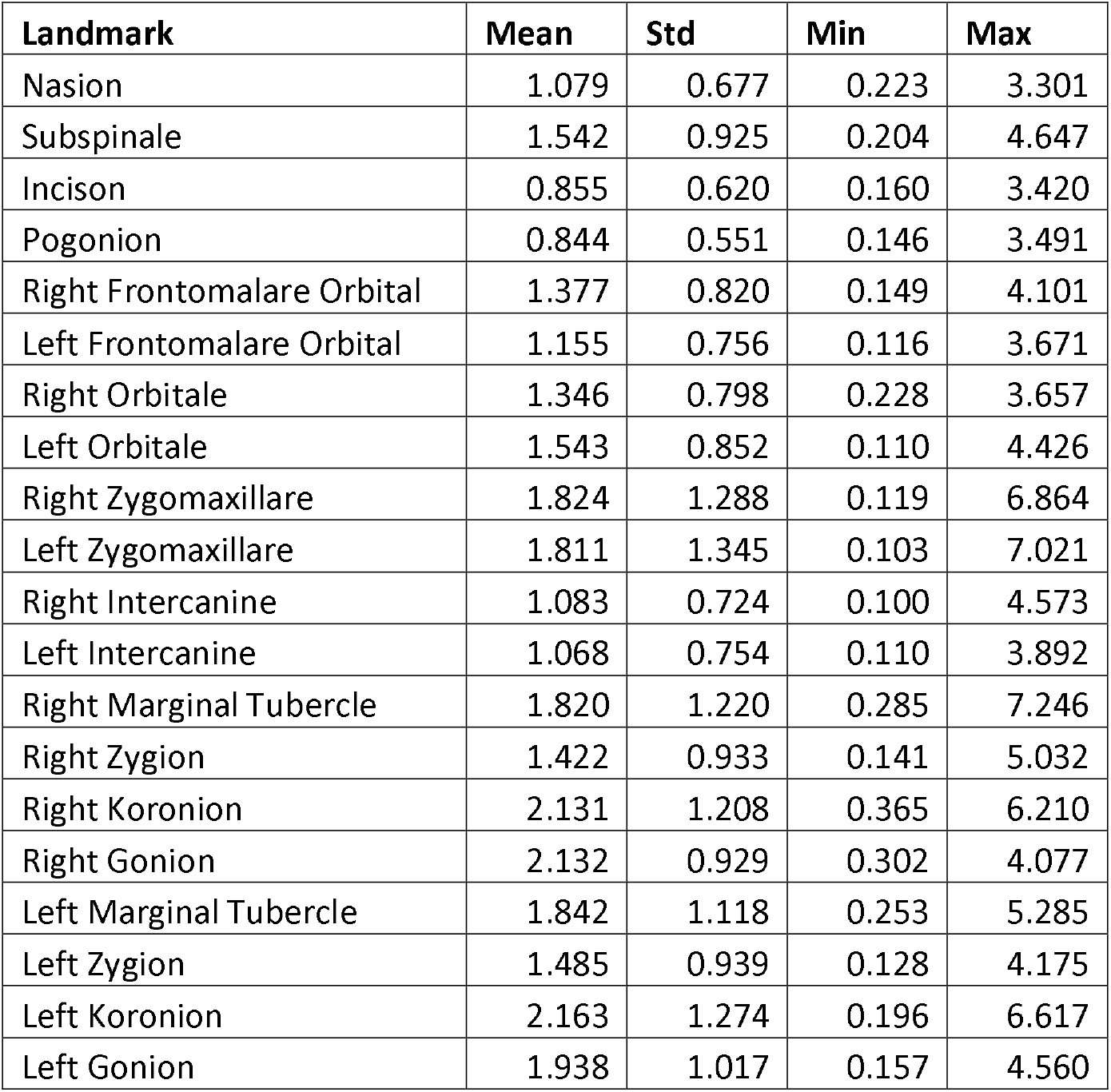
Descriptive statistics for the Euclidean distances between the average manual and average automatic landmarks in mm.

**Table 3:** ANOVA on centroid size of manual and automatic landmark configurations. Skull and method were inputted as factors.

|  | Df | Sum Sq | Mean Sq | F value | Pr(>F) |
| --- | --- | --- | --- | --- | --- |
| **Skull** | 30 | 22825.48 | 760.849 | 958.120 | <0.001 |
| **Observer** | 2 | 241.90 | 120.951 | 152.311 | <0.001 |
| **Method** | 1 | 2.84 | 2.844 | 3.581 | 0.062 |
| **Skull\*Observer** | 60 | 20.13 | 0.336 | 0.423 | 1.000 |
| **Residuals** | 92 | 73.06 | 0.794 |  |  |

**Figure 3:**
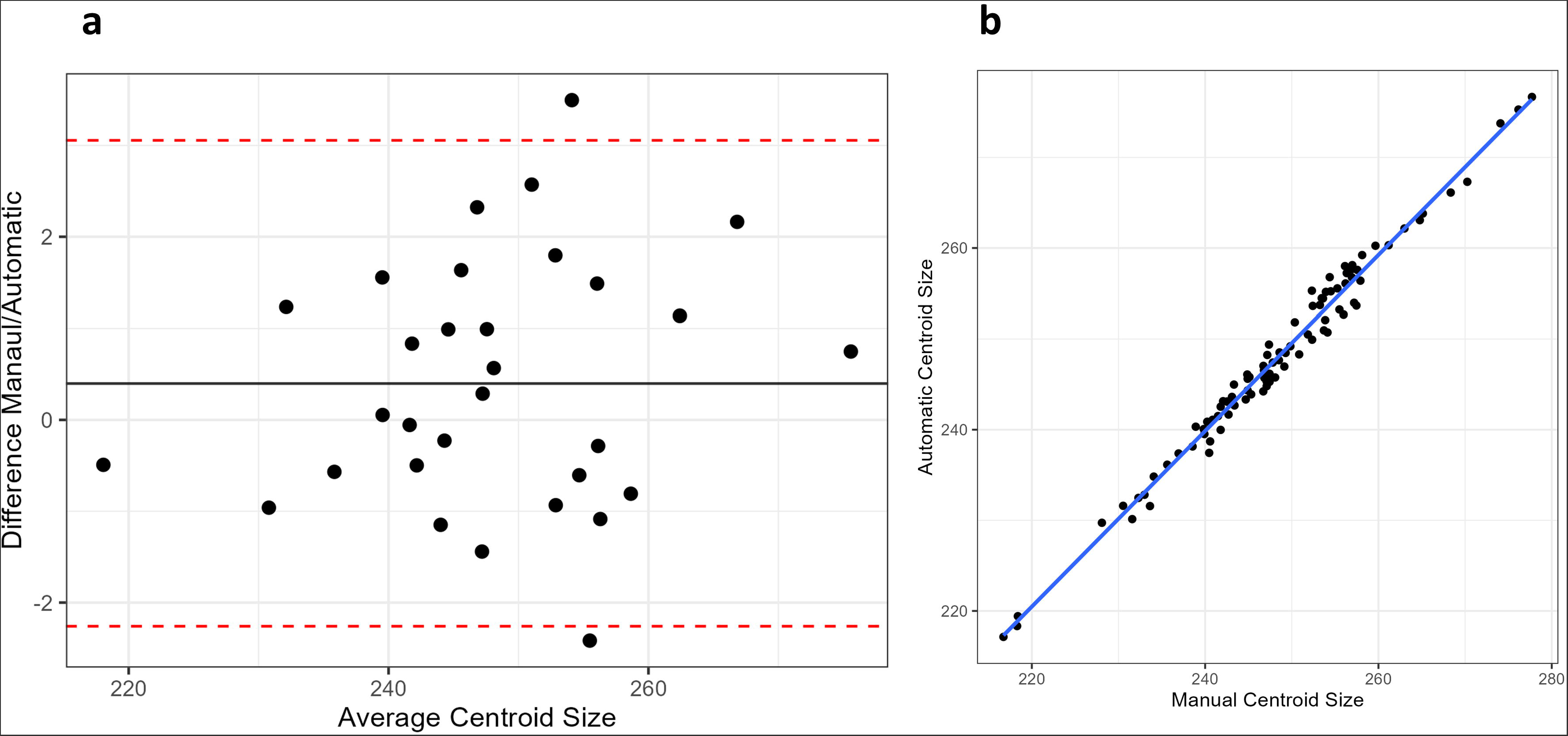
A: Bland-Altman showing agreement of centroid size between manual and automatic landmark configurations, B: Comparison of manual and automatic centroid sizes.

**Figure 4:**
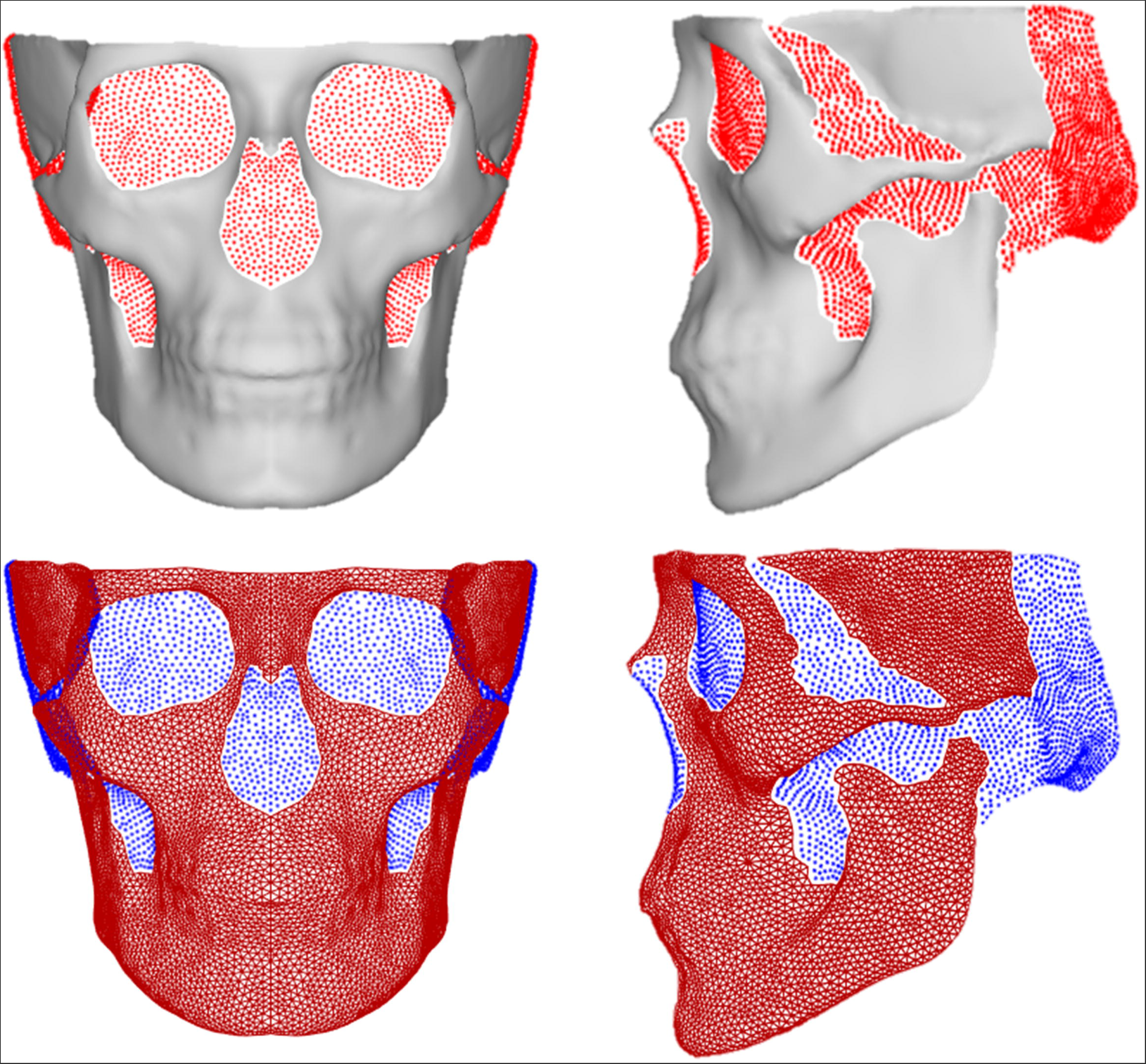
Visual representation of the “true” stable landmarks and the “gap” landmarks. Vertex IDs for these landmarks can be found in the supplementary file.

An ANOVA over the centroid sizes with observer, skull and nested observer/iteration as factors showed that all factors contributed significantly to variation in centroid size (Supplementary Table S4). A MANOVA was performed on the GPA aligned manual landmarks with observer, skull, and nested observer/iteration as predictors (Supplementary Table S5). The skull itself contributed the largest amount of variation (R^2^=94%) while the other predictors did not contribute.

By treating the automatic landmarks obtained by using each observer’s manual landmarks as the “gold standard’ we could calculate inter-observer error for the automatic landmarks. The mean standard deviation was smaller for automatic landmarks (0.77mm) than for manual landmarks (0.902mm). Levene’s test for equal variance also showed that the variation within the automatic landmarks was significantly smaller than that in the manual landmarks (Supplementary Table S6). The standard deviation of the average over the xyz axes was smaller for the automatic landmarks than for the manual landmarks for all landmarks, and with less extreme outliers (Supplementary Figure S7). 80% of variation within the automatic landmarks was explained by individual variation, while 19% was explained by observer differences (Supplementary Table S5).

### ACCURACY

#### Euclidean Distance Comparison

As a first measure of accuracy, the Euclidean distance between the manual and automatic landmarks was calculated (Table 2). The average distance over all landmarks was 1.5mm, with a range from 0.1mm (Intercanine) to 7.2mm (Marginal Tubercle). A Bland-Altman plot was generated to evaluate if specific axes show higher discordance (Supplementary Figure 7). The variation shown on the principal axes found in Supplementary Figure 8 illustrates that often the first axis follows the contour of the skull. Landmarks with clear definition points in all axes show smaller errors while those that have a sliding placement along a contour generate larger errors. ICC for each axes are 0.998 or above showing high agreement.

#### Centroid Size Comparison

A Bland-Altman plot shows a mean difference in centroid size of 0.4mm between the two methods, in addition to high concordance between centroid sizes over all the CBCT skulls (Figure 3). ICC calculated from centroid sizes was 0.99 (95% CI: 0.986 < ICC < 0.994) showing negligible differences in the landmarking method. An ANOVA on centroid size with skull and method as factors showed that the method itself was not significant (p=0.06), while skull and observer were highly significant (p<0.001) (Table 3).

#### Shape Comparison

A MANOVA performed on the GPA aligned coordinates with skull, observer, and method as predictors showed that the Method could only explain 2% of the variation, while observer accounted for 4.6% and the skulls were 40% (Supplementary Table S7). In addition, a MANOVA was performed on the first 13 PCs (explaining ∼95% of the variation) of the auto and manual landmarks using the same predictors. Only observer and skull significantly affected PC scores, the method was not significant (Supplementary Table S8).

### ESTABLISHED LANDMARKS & SUCCESSFUL IMAGE APPLICATION

Of the 9,999 original quasi-landmarks, 6,707 were defined as “true” landmarks pertaining to locations on the physical skull. Those landmarks that fill gaps were flagged and can visually be seen to locate to 1) gaps such as the eye sockets and nose, 2) areas near the back of the skull that are not well represented in CBCT images (Figure 4). A basic analysis of sexual dimorphism and PCA was performed on all 9,999 landmarks and can be found in the Supplementary Material (GIF 1 (Sex) and 2 (PC1)) as well as our website; https://walshlab.sitehost.iu.edu/pages/craniofacial.html. The CT image taken from the MUG500+ dataset [36] was also successfully masked (Supplementary Figure S7).

## DISCUSSION

While our understanding of human facial morphology and its variation has made notable progress, particularly with regards soft tissue variation, our knowledge of the underlying hard tissue structures has significantly trailed. This disparity can be primarily attributed to the paucity of extensive skull datasets and the limited development of advanced 3D morphometric methods for skull shape analysis. To overcome this, the utilization of CBCT imagery, typically taken by dentists provides a more accessible solution whilst offering superior resolution in the form of skull mesh reconstructions. Although the analysis of CBCT-derived skull data is not devoid of inherent challenges; they frequently exhibit only partial cranial representations and are characterized by the absence of posterior and superior cranial segments, alongside potential holes in the mesh structure. A consequence of this is that typically 3D masks must be adjusted to accommodate these challenging structural limitations. In addition, the meshes are often irregular, and prone to artifacts stemming from minor movements, dental interventions, and the utilization of head supports during imaging procedures. Lastly, CBCT skull meshes incorporate all hard tissue, including structures within the skull, making these meshes complex and difficult to mask. CT images show similar issues: while the mesh is often of the complete skull, artefacts are more extreme, and the surface texture is more irregular.

With the aim of enhancing the manageability of hard tissue scans, we designed and provide a craniofacial template bone mask that not only reduces the complexity of skull scans by targeting a specific area of interest but also leverages a previously proven 3D phenotyping methodology [27]. Our craniofacial template bone mask encompasses 9,999 quasi-landmarks focusing solely on the externally visible aspect of the cranial structure, of which 6,707 define points on the physical skull. Through the process of shrink-wrapping and subsequent reduction of the skull meshes derived from Cone Beam Computed Tomography (CBCT) and Computed Tomography (CT) imagery, we achieve a substantial reduction in polygon count, facilitating more manageable data handling, particularly on less robust computational platforms. This innovative approach of wrapping the skulls also permits Meshmonk to mask the skull without the need to navigate complex interior structures or address gaps in the mesh. While the use of meshmonk on a full skull has been published [31], this application could not be applied to our CBCT or CT scans as the template is not freely available. The authors also note that the mask consisted of ∼177,000 quasi-landmarks and necessitated the use of the full skull captured using the same imaging modality.

This study aimed to provide an alternative craniofacial bone template that has been fully evaluated with regards to its reliability and accuracy. Meshmonk requires a few manually placed landmarks for its preliminary registration process and our findings reveal that variations in the placement of these landmarks do introduce some degree of error in the masking process. However, these errors predominantly manifest along the periphery of the mask and correspond to regions that may not always be entirely captured in CBCT scans. Specifically, the absence of posterior segments can result in the mask coalescing in this region. This is especially visible when calculating the “true” landmarks as it was evident these posterior points were not well represented in our cohort and thus flagged as “gaps”. Consequently, it is plausible that this issue may be more attributable to limitations inherent to the CBCT imaging technique rather than deficiencies in the masking methodology. Notably, the maximum Meshmonk error is higher than that seen for a mandible mask [30], albeit confined to these specific regions. Our errors were reduced by modifying the Meshmonk settings “NRM.FlagFloatingBoundary” and “NRM.FlagTargetBoundary” (for CT both were set to true, for CBCT only the first was set to true) and recommend that these settings be tested in combination with our script to produce a type of ‘quality control’ test image that depicts the mask overlaying the original mesh (see Supplementary File) to define the best settings for each user’s imagery. Enhanced CBCT image preprocessing techniques and the acquisition of more comprehensive CBCT images would potentially ameliorate this error.

We also considered the intra- and inter-observer errors associated with manual landmarking of the skull. 20 landmarks were selected that were easily identifiable on the skull. Only one of the observers was trained in cranial landmarking, resulting in a significantly lower overall RMS error (0.66mm) in comparison to the other two observers (1.11mm and 0.94mm). These values are similar to those reported in other studies [27, 30, 31], albeit slightly higher than findings from a study employing specialized landmarking software at some landmarks [31]. As expected, the inter-observer error was higher (1.44mm). When this was compared to the automatic landmarking error over the 3 different iterations of initial landmarks, the error was more than 6x smaller (0.12mm).

Automatic phenotyping demonstrated good accuracy when compared to manual landmarking. To determine the corresponding automatic landmarks, the manual landmarks served as the gold standard. To mitigate bias, a leave-one-out approach was used. Analysis revealed a variation in Euclidean distance between manual and automatic landmarks ranging from 0.10mm - 7.24mm, with an average of 1.5mm. Notably, landmarks exhibiting higher values were often associated with regions that were less effectively masked (e.g., gonion where the mask did not always align with the edge of the skull) and were prone to higher inter-observer errors, as observed in previous studies [30]. Previous research has reported comparable errors between manual and automatic landmark placements ranging from 2.19mm [31], 1.4mm [30], 2.01mm [42] and 1.26mm [27]. And our ICC values consistently demonstrated high levels of agreement (ICC > 0.9). Interestingly, observer-related variability contributed to 19% of the variation in automatic landmark placement, but this factor was negligible in the case of manual placement. This observation suggests that variations introduced by different observers had an impact on the calculations for automatic landmark placement, as the manual placements were used to calculate the automatic landmark placement. Notably, a considerably smaller proportion of variation remained unexplained for the automatic landmarks (0.4% vs. 6%). An ANOVA on centroid size and a Multivariate Analysis of Variance (MANOVA) conducted on the first 13 Principal Components (PCs) demonstrated clearly that the method employed did not exert a significant influence on the variation observed in landmark configurations or shape.

The conventional use of manual landmarks as the “gold standard,” has been applied and observed in previous studies [27, 30, 43], it also raises certain inherent issues. Specifically in this case, two out of the three observers lacked prior training in landmarking. This introduces more variability in the automatic landmark placement. Moreover, our analysis focused on a limited set of 20 landmarks, therefore the accuracy of the remaining quasi-landmarks was not systematically assessed. However, given the low error associated with automatic landmarking, it is reasonable to assume that these additional landmarks exhibit a comparable level of accuracy, which remain significantly lower than manual landmarking errors. Another limiting factor was the quality of the CBCT images. Despite our efforts to select images with minimal artifacts, many still exhibited minor missing portions and common CBCT imaging artifacts. Additionally, there was considerable variability in the extent of image coverage, particularly in the posterior regions of the scans due to the CBCT machines’ limited FOV. Thus, it was expected that our results showed higher errors than commonly seen when landmarking complete 3D skull scans or physical skulls. However, these types of scans represent the reality of present data sets. It also underscores the viability of the shrink-wrapped craniofacial bone mask as an effective method that more easily facilitates the comparison of the human skulls geometric shape, particularly when derived from CBCT and CT imagery. To visualize the potential use of our mask on these types of scans, we provide a PCA and sexual dimorphism analysis that could easily show morphological variation even within our small sample set. Many research groups working on geometric morphometrics and the genetics behind skull and face shape may lack formal anthropological training, leading to elevated manual landmarking errors akin to those observed with observers 2 and 3. By providing a method for stable automatic landmarking, this error is minimized. It also reduces time investment, enhances objectivity, and has the capacity to analyze a greater number of landmarks. Additionally, the deliberate focus on the external aspects of the skull and the creation of a single-plane craniofacial bone mask not only reduces computational resource requirements but also standardizes subsequent analyses. We have also supplied a comprehensive workflow for the shrink-wrapping procedure, quality control scripts, and the vertex IDs for the stable landmarks, rendering this method easily adaptable in research laboratories without the need for specialized training. The adoption of a standardized mask further facilitates the efficient comparison of data across various studies. While this craniofacial bone mask may only encompass part of the cranium, this allows it to be applicable to CBCTs collected on devices with limited focal views. In the future we plan to extend this work to a full cranial mask and utilize the development of superior CT and CBCT scanners [44, 45] and deep learning for DICOM segmentation [46]. Ultimately, we envision that our work will pave the way for genetic association studies pertaining to cranial shape and high-resolution investigations into the genetic determinants influencing craniofacial bone morphology.

## CONCLUSION

Within this study we designed and provide a freely available 3D craniofacial template bone mask for the dense 3D phenotyping of skull meshes exported from CBCT/CT scans in addition to a tutorial outlining the procedure for preparing these images for masking. The provided template can be used within the Meshmonk framework, facilitating the generation of high-density cranial landmarks for subsequent analyses with minimal manual intervention and in a notably efficient manner. Our methodology has demonstrated a high level of accuracy, with substantially reduced errors when compared to manual landmark placement. This standardized approach not only enhances reliability and precision but also minimizes the potential for errors in landmark identification and placement in hard tissue structures of the human face.

## ACKNOWLEDGEMENTS

We thank all the individuals and families that gave their time and took part in these facial studies. We would also like to thank Dr. Kelton Stewart and Brenda McClarnon at IU School of Dentistry for facilitating and taking the CBCT images used in this study. IUI personnel, data collection, and analyses were supported by the National Institute of Health (NIDCR 1R15DE031929).

## AUTHOR CONTRIBUTIONS

FW: Conceived and designed the analyses, collected the data, performed the analyses, wrote the manuscript

NH: Collected the data, performed the analyses

ND: Performed the analyses

HM: Contributed data or analyses tools

PC: Contributed data or analyses tools

SW: Collected the data, conceived, designed the analyses, revised the manuscript & oversaw the project

## DATA AVAILABILITY

The craniofacial template bone mask that is the basis of this work can be downloaded from the supplementary file and our website; https://walshlab.sitehost.iu.edu/pages/craniofacial.html

The MeshMonk (v.0.0.6) spatially dense facial-mapping software is provided by KU Leuven and is free to use for academic purposes (https://gitlab.kuleuven.be/mirc/meshmonk). All code used for analyses was modified from a previous publication (https://doi.org/10.1038/s41588-020-00741-7). Further underlying code is available as part of the Matlab software (https://www.mathworks.com) as well as 3D Slicer (https://www.slicer.org/), Blender (https://www.blender.org/), Meshmixer (https://meshmixer.com/) and MeVisLab (https://www.mevislab.de/).

