## Supplementary Tables & Figures for "Automated 3D Landmarking of the Skull: A Novel Approach for Craniofacial Analysis"

**TITLE**

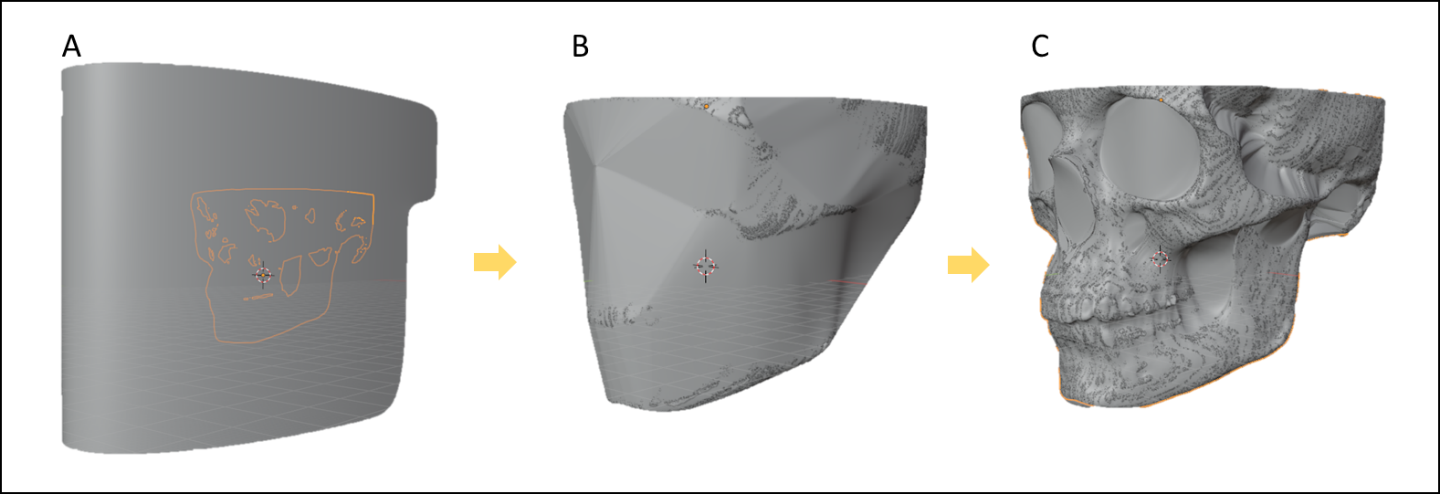
**Supplementary Figure S1:** Shrink-wrapping in Blender. A: a deformed half cylinder is placed around the skull, B: using Blender’s shrinkwrapping tool the cylinder is wrapped to the skull, C: This step is repeated 6x while applying the subdivision surface modified in between rounds.

**Supplementary Figure**
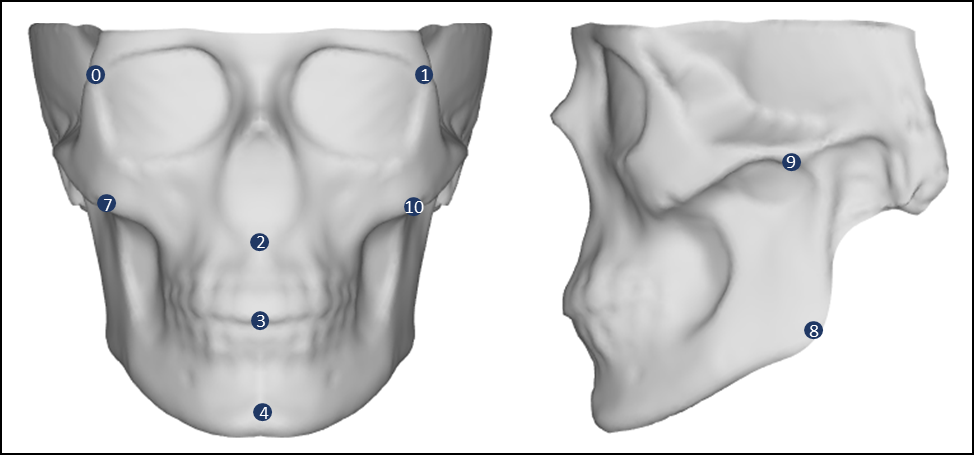
**S2:** Landmarks used for Meshmonk initialization.

| **Landmark** | **O1** | **O2** | **O3** | **IO** |
| --- | --- | --- | --- | --- |
| Nasion | 0.569 | 0.727 | 0.897 | 1.610 |
| Subspinale | 0.657 | 0.812 | 0.903 | 1.498 |
| Incison | 0.410 | 0.597 | 0.656 | 0.920 |
| Pogonion | 0.459 | 0.730 | 0.658 | 1.788 |
| Right Frontomalare Orbital | 0.483 | 0.734 | 0.790 | 1.464 |
| Left Frontomalare Orbital | 0.531 | 0.836 | 0.703 | 1.977 |
| Right Orbitale | 0.806 | 0.984 | 1.234 | 1.792 |
| Left Orbitale | 0.918 | 0.899 | 0.918 | 1.653 |
| Right Zygomaxillare | 1.106 | 2.056 | 1.388 | 2.393 |
| Left Zygomaxillare | 0.980 | 1.805 | 1.494 | 2.053 |
| Right Intercanine | 0.684 | 1.076 | 1.182 | 1.448 |
| Left Intercanine | 0.688 | 0.911 | 1.202 | 1.311 |
| Right Marginal Tubercle | 0.455 | 1.123 | 0.627 | 0.880 |
| Right Zygion | 0.779 | 1.603 | 1.080 | 1.948 |
| Right Koronion | 0.600 | 1.058 | 0.816 | 0.738 |
| Right Gonion | 0.577 | 1.071 | 0.723 | 0.914 |
| Left Marginal Tubercle | 0.511 | 1.193 | 0.695 | 0.946 |
| Left Zygion | 0.662 | 1.369 | 1.344 | 1.598 |
| Left Koronion | 0.634 | 1.157 | 0.812 | 0.850 |
| Left Gonion | 0.730 | 1.434 | 0.746 | 1.069 |
| Mean | 0.662 | 1.109 | 0.943 | 1.442 |
| Std | 0.180 | 0.372 | 0.267 | 0.462 |
| Min | 0.410 | 0.597 | 0.627 | 0.738 |
| Max | 1.106 | 2.056 | 1.494 | 2.393 |

**Supplementary Table S1:** Intra-Observer (O1-O3) and Inter-Oberver (IO) error shown as the RMS distance (in mm) to the centroid over the 3 landmarking rounds/three observers per landmark.

**Supplementary Table S2:** RMS distance (in mm) to centroid over the three meshmonk interations for the 20 landmarks

| **Landmark** | **RMS (mm)** |
| --- | --- |
| Nasion | 0.052 |
| Subspinale | 0.290 |
| Incison | 0.115 |
| Pogonion | 0.077 |
| Right Frontomalare Orbital | 0.092 |
| Left Frontomalare Orbital | 0.079 |
| Right Orbitale | 0.122 |
| Left Orbitale | 0.076 |
| Right Zygomaxillare | 0.122 |
| Left Zygomaxillare | 0.124 |
| Right Intercanine | 0.086 |
| Left Intercanine | 0.095 |
| Right Marginal Tubercle | 0.059 |
| Right Zygion | 0.080 |
| Right Koronion | 0.058 |
| Right Gonion | 0.322 |
| Left Marginal Tubercle | 0.065 |
| Left Zygion | 0.068 |
| Left Koronion | 0.052 |
| Left Gonion | 0.347 |
| Mean | 0.119 |
| Std | 0.086 |
| Min | 0.052 |
| Max | 0.347 |

**Supplementary Figure S3:** RMS distance (in mm) to centroid over the three meshmonk iterations for all 9,999 quasi-landmarks.

|  | **Mean** | **95% CI Mean** | | **SD** | **Min** | **Max** |
| --- | --- | --- | --- | --- | --- | --- |
| **Automated** | 0.1190 | 0.0769 | 0.1611 | 0.0899 | 0.0519 | 0.3468 |

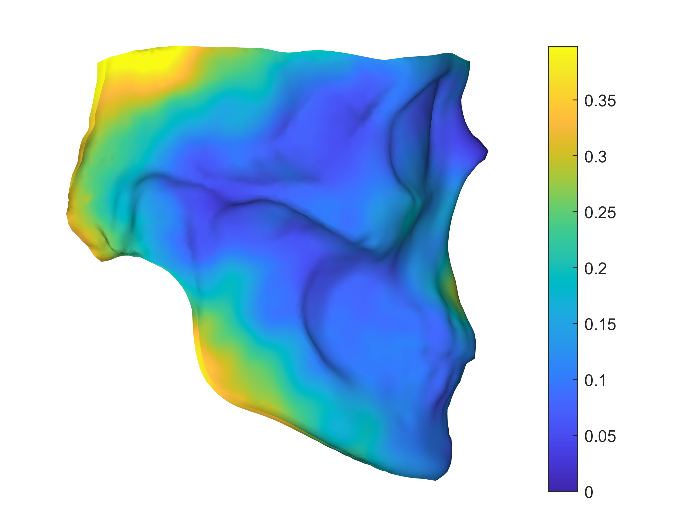

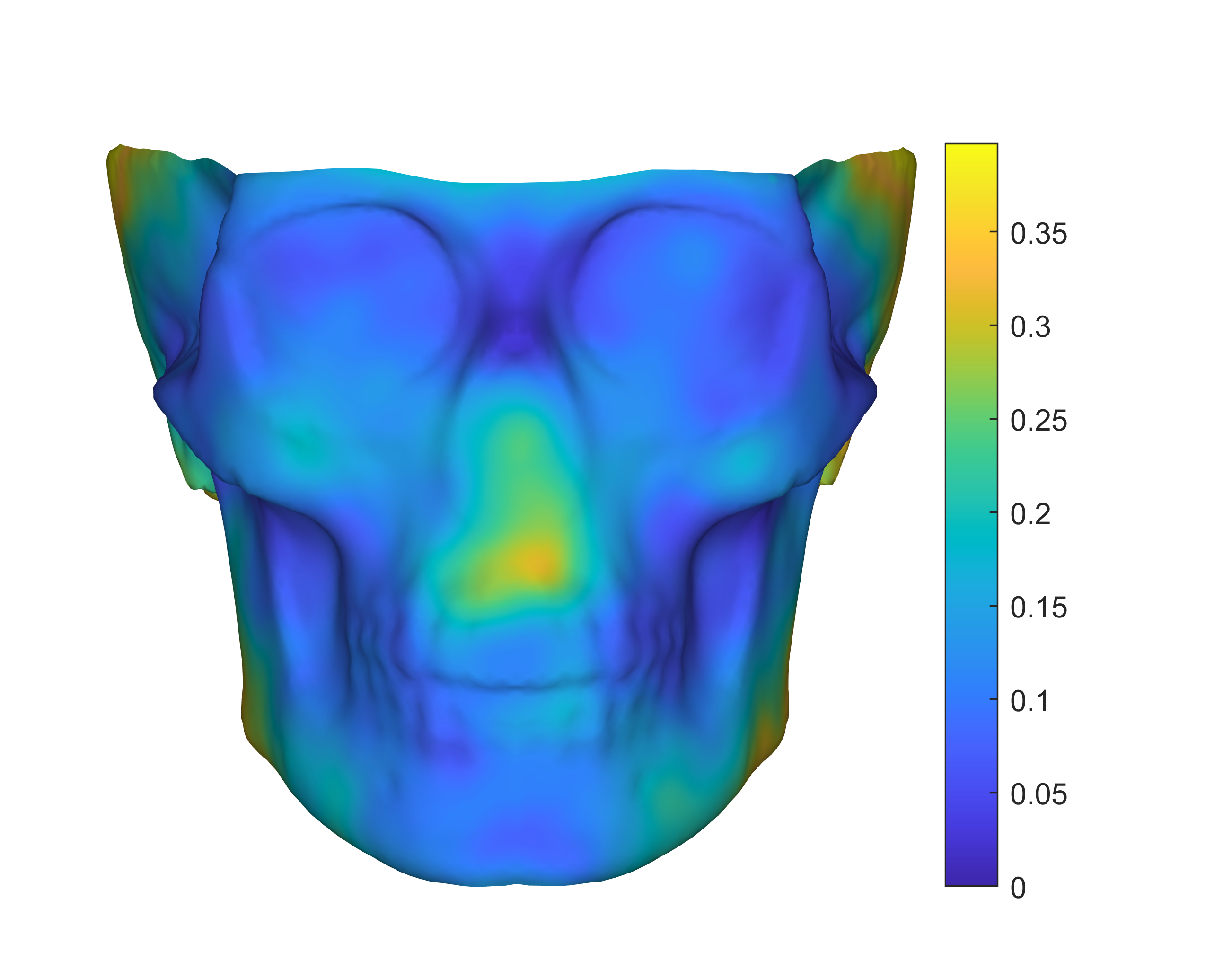

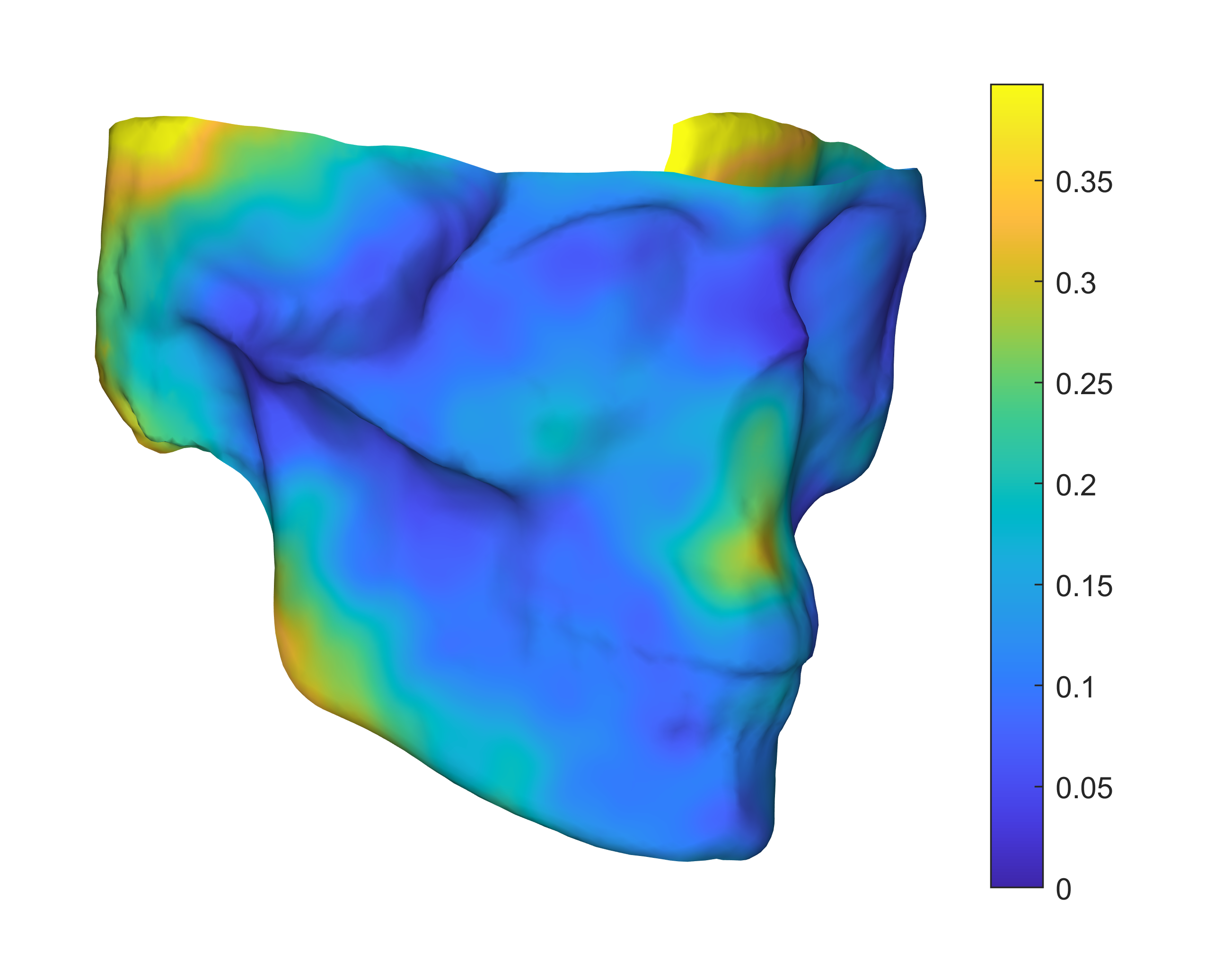

| **Landmark** | **Red** | **Blue** | **Green** |
| --- | --- | --- | --- |
| Nasion | 0.808 | 0.215 | 0.151 |
| Subspinale | 0.801 | 0.351 | 0.158 |
| Incison | 0.487 | 0.367 | 0.225 |
| Pogonion | 0.629 | 0.291 | 0.086 |
| Right Frontomalare Orbital | 0.670 | 0.352 | 0.103 |
| Left Frontomalare Orbital | 0.664 | 0.402 | 0.145 |
| Right Orbitale | 1.032 | 0.483 | 0.182 |
| Left Orbitale | 0.889 | 0.423 | 0.160 |
| Right Zygomaxillare | 1.806 | 0.780 | 0.379 |
| Left Zygomaxillare | 1.598 | 0.794 | 0.389 |
| Right Intercanine | 1.151 | 0.409 | 0.172 |
| Left Intercanine | 1.072 | 0.373 | 0.153 |
| Right Marginal Tubercle | 0.995 | 0.427 | 0.186 |
| Right Zygion | 1.200 | 0.663 | 0.230 |
| Right Koronion | 0.753 | 0.576 | 0.172 |
| Right Gonion | 0.827 | 0.409 | 0.144 |
| Left Marginal Tubercle | 0.867 | 0.527 | 0.222 |
| Left Zygion | 1.283 | 0.569 | 0.188 |
| Left Koronion | 0.820 | 0.679 | 0.250 |
| Left Gonion | 1.082 | 0.375 | 0.114 |

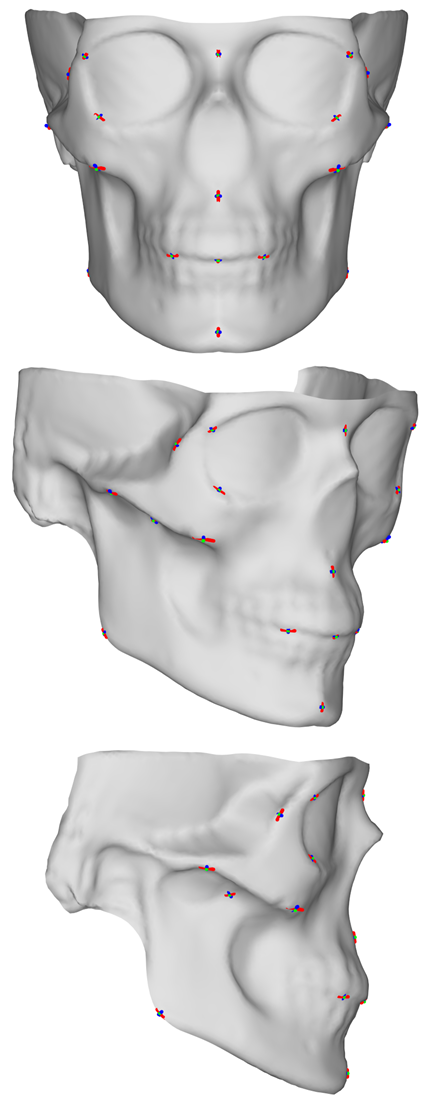
**Supplementary Figure S4:** Visual and tabular variation of the intra-observer error over the 20 manually placed landmarks. Variation is shown as the three largest dimensions and in which direction this variation occured.

**Supplementary Figure S5:** Visual and tabular variation of the inter-observer error over the 20 manually placed landmarks. Variation is shown as the three largest dimensions and in which direction this variation occured.

| **Landmark** | **Red** | **Blue** | **Green** |
| --- | --- | --- | --- |
| Nasion | 1.677 | 0.214 | 0.182 |
| Subspinale | 1.531 | 0.309 | 0.231 |
| Incison | 0.883 | 0.379 | 0.179 |
| Pogonion | 1.816 | 0.278 | 0.269 |
| Right Frontomalare Orbital | 1.433 | 0.478 | 0.201 |
| Left Frontomalare Orbital | 1.906 | 0.659 | 0.266 |
| Right Orbitale | 1.581 | 0.861 | 0.279 |
| Left Orbitale | 1.427 | 0.858 | 0.213 |
| Right Zygomaxillare | 2.104 | 1.239 | 0.370 |
| Left Zygomaxillare | 1.737 | 1.323 | 0.340 |
| Right Intercanine | 1.452 | 0.446 | 0.194 |
| Left Intercanine | 1.318 | 0.540 | 0.147 |
| Right Marginal Tubercle | 0.919 | 0.515 | 0.148 |
| Right Zygion | 1.514 | 1.359 | 0.343 |
| Right Koronion | 0.723 | 0.443 | 0.145 |
| Right Gonion | 0.928 | 0.250 | 0.114 |
| Left Marginal Tubercle | 0.972 | 0.379 | 0.169 |
| Left Zygion | 1.260 | 1.076 | 0.268 |
| Left Koronion | 0.804 | 0.466 | 0.206 |
| Left Gonion | 1.103 | 0.247 | 0.132 |

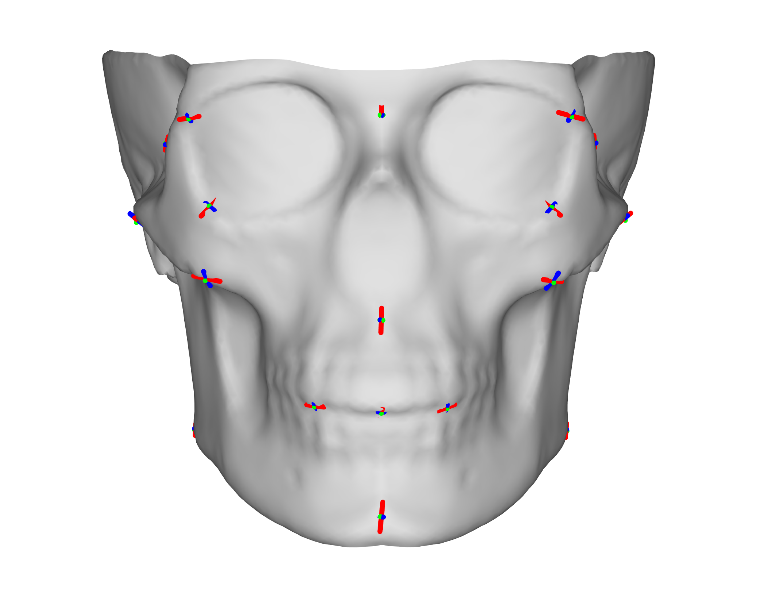

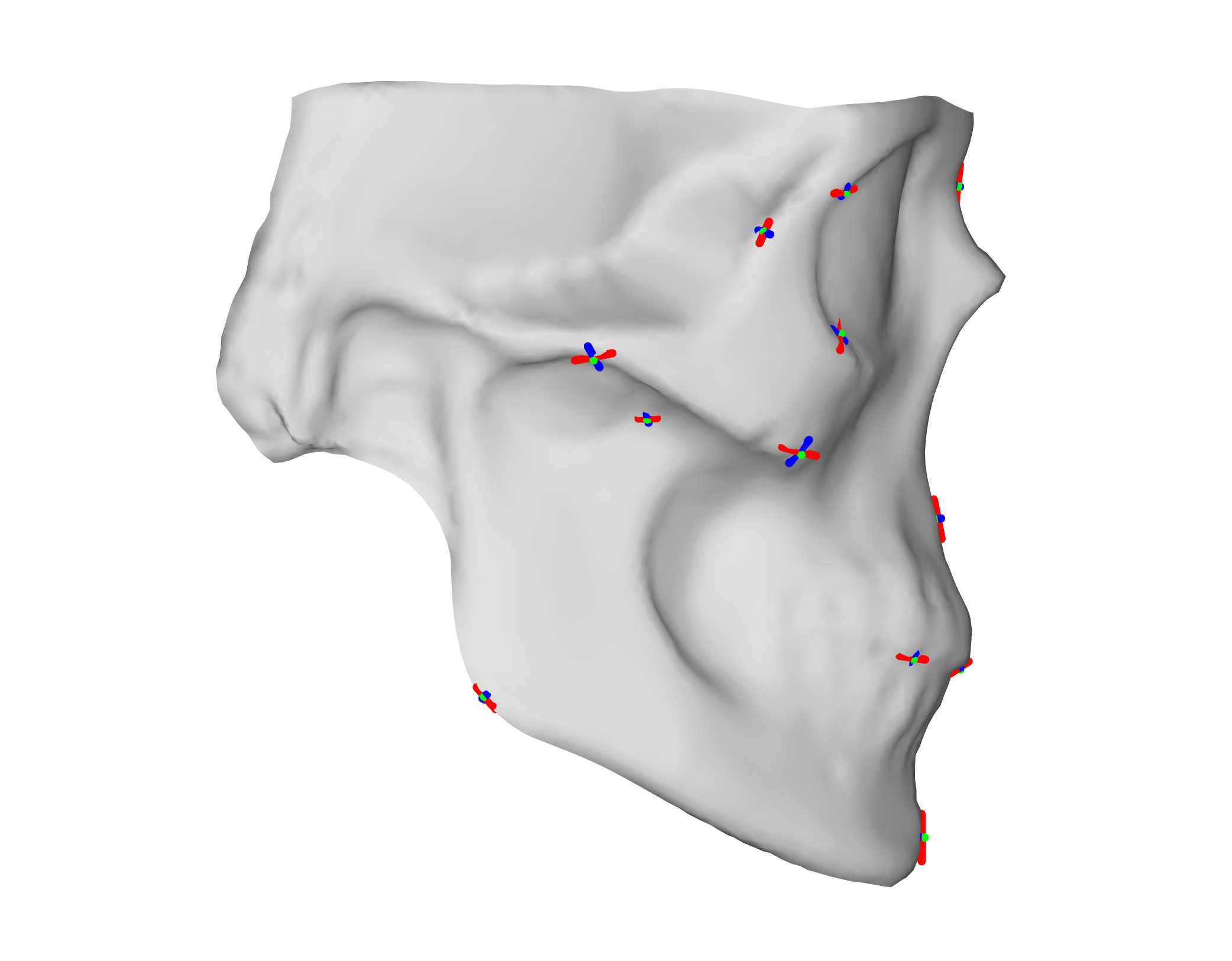

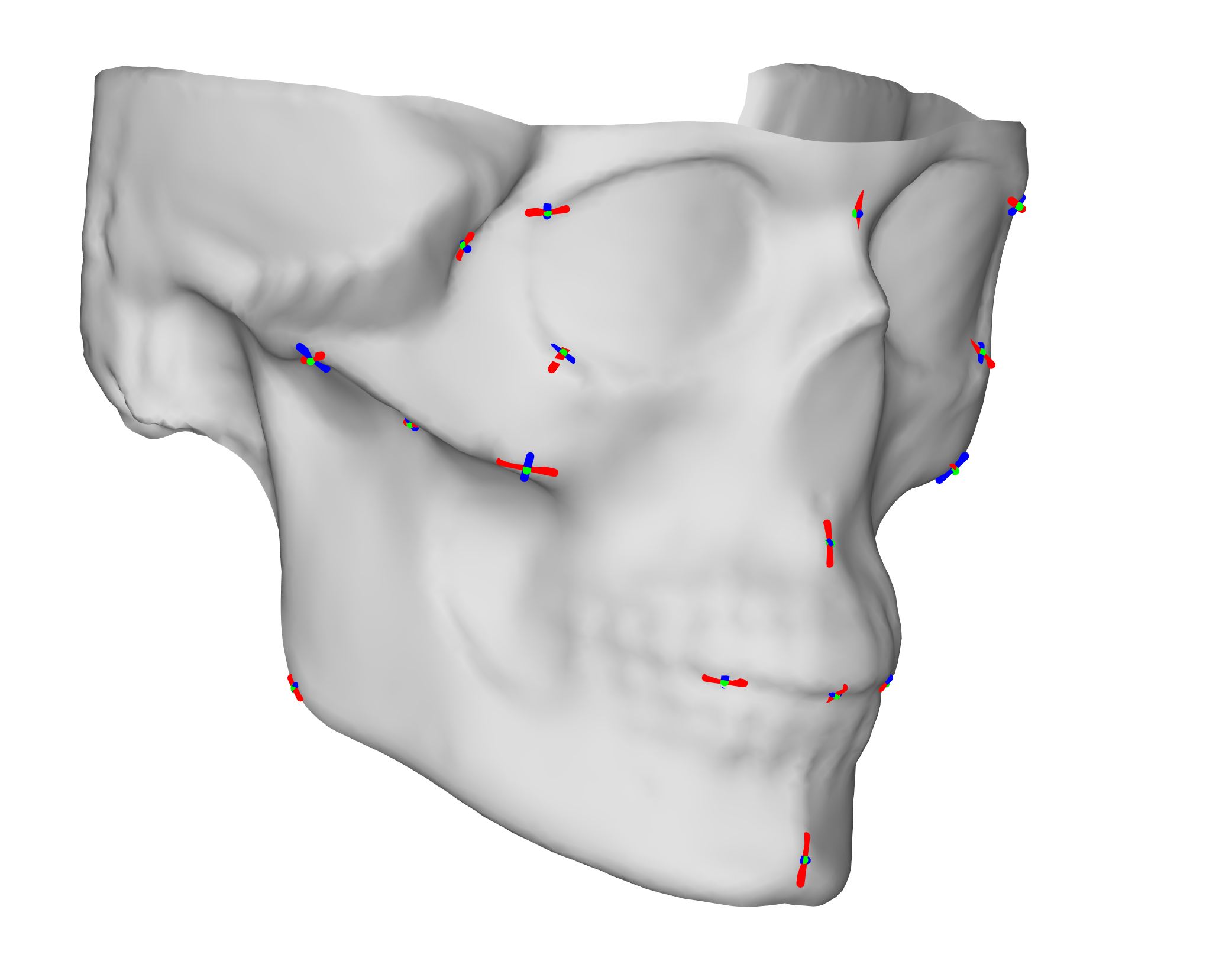

**Supplementary Table S3:** Intra-observer error shown as the standard deviation of the landmarks along the x, y, and z axes per landmark. Values are the standard deviation in mm.

|  | **Observer 1** | | | | **Observer 2** | | | | **Observer 3** | | | |
| --- | --- | --- | --- | --- | --- | --- | --- | --- | --- | --- | --- | --- |
| **Landmark** | **x** | **y** | **z** | **Mean** | **x** | **y** | **z** | **Mean** | **x** | **y** | **z** | **Mean** |
| **Nasion** | 0.190 | 0.656 | 0.055 | 0.300 | 0.259 | 0.797 | 0.169 | 0.408 | 0.256 | 1.030 | 0.166 | 0.484 |
| **Subspinale** | 0.828 | 0.543 | 0.556 | 0.642 | 1.357 | 0.570 | 1.438 | 1.122 | 1.279 | 0.839 | 0.844 | 0.987 |
| **Incison** | 0.486 | 0.321 | 0.564 | 0.457 | 0.704 | 0.480 | 0.883 | 0.689 | 1.051 | 0.352 | 0.870 | 0.758 |
| **Pogonion** | 0.516 | 0.252 | 0.569 | 0.446 | 0.634 | 0.435 | 0.712 | 0.594 | 1.035 | 0.365 | 0.931 | 0.777 |
| **Right Frontomalare Orbital** | 0.080 | 0.431 | 0.291 | 0.267 | 0.434 | 0.994 | 0.735 | 0.721 | 0.125 | 0.628 | 0.369 | 0.374 |
| **Left Frontomalare Orbital** | 0.256 | 0.234 | 0.852 | 0.447 | 1.081 | 0.461 | 1.407 | 0.983 | 0.207 | 0.297 | 1.244 | 0.583 |
| **Right Orbitale** | 0.350 | 0.380 | 0.451 | 0.394 | 0.581 | 0.557 | 0.925 | 0.688 | 0.428 | 0.453 | 0.715 | 0.532 |
| **Left Orbitale** | 0.152 | 0.446 | 0.492 | 0.364 | 0.295 | 0.915 | 0.817 | 0.676 | 0.205 | 0.503 | 0.624 | 0.444 |
| **Right Zygomaxillare** | 0.127 | 0.456 | 0.365 | 0.316 | 0.557 | 0.973 | 0.721 | 0.750 | 0.155 | 0.659 | 0.440 | 0.418 |
| **Left Zygomaxillare** | 0.225 | 0.198 | 0.718 | 0.380 | 0.858 | 0.375 | 1.243 | 0.825 | 0.213 | 0.319 | 1.572 | 0.701 |
| **Right Intercanine** | 0.297 | 0.367 | 0.536 | 0.400 | 0.772 | 0.624 | 0.873 | 0.756 | 0.387 | 0.525 | 0.658 | 0.524 |
| **Left Intercanine** | 0.284 | 0.687 | 0.189 | 0.387 | 0.425 | 0.773 | 0.267 | 0.488 | 0.380 | 0.964 | 0.240 | 0.528 |
| **Right Marginal Tubercle** | 0.123 | 0.623 | 0.588 | 0.445 | 0.258 | 1.129 | 1.210 | 0.866 | 0.149 | 0.605 | 0.624 | 0.459 |
| **Right Zygion** | 0.338 | 0.255 | 0.180 | 0.258 | 0.330 | 0.443 | 0.383 | 0.385 | 0.429 | 0.403 | 0.405 | 0.412 |
| **Right Koronion** | 0.290 | 0.444 | 0.071 | 0.268 | 0.335 | 0.789 | 0.110 | 0.411 | 0.296 | 0.685 | 0.151 | 0.377 |
| **Right Gonion** | 0.296 | 0.302 | 0.361 | 0.320 | 0.451 | 0.505 | 0.491 | 0.482 | 0.378 | 0.548 | 0.649 | 0.525 |
| **Left Marginal Tubercle** | 0.370 | 0.328 | 0.348 | 0.349 | 0.543 | 0.584 | 0.553 | 0.560 | 0.397 | 0.528 | 0.449 | 0.458 |
| **Left Zygion** | 0.685 | 0.462 | 0.449 | 0.532 | 0.807 | 0.634 | 0.532 | 0.658 | 0.879 | 0.796 | 0.825 | 0.833 |
| **Left Koronion** | 0.805 | 0.480 | 0.532 | 0.606 | 0.699 | 0.622 | 0.482 | 0.601 | 0.690 | 0.567 | 0.619 | 0.625 |
| **Left Gonion** | 0.829 | 0.437 | 0.882 | 0.716 | 1.391 | 0.541 | 1.820 | 1.251 | 1.258 | 0.573 | 0.857 | 0.896 |
| **Mean** | 0.376 | 0.415 | 0.452 | 0.415 | 0.639 | 0.660 | 0.789 | 0.696 | 0.510 | 0.582 | 0.663 | 0.585 |

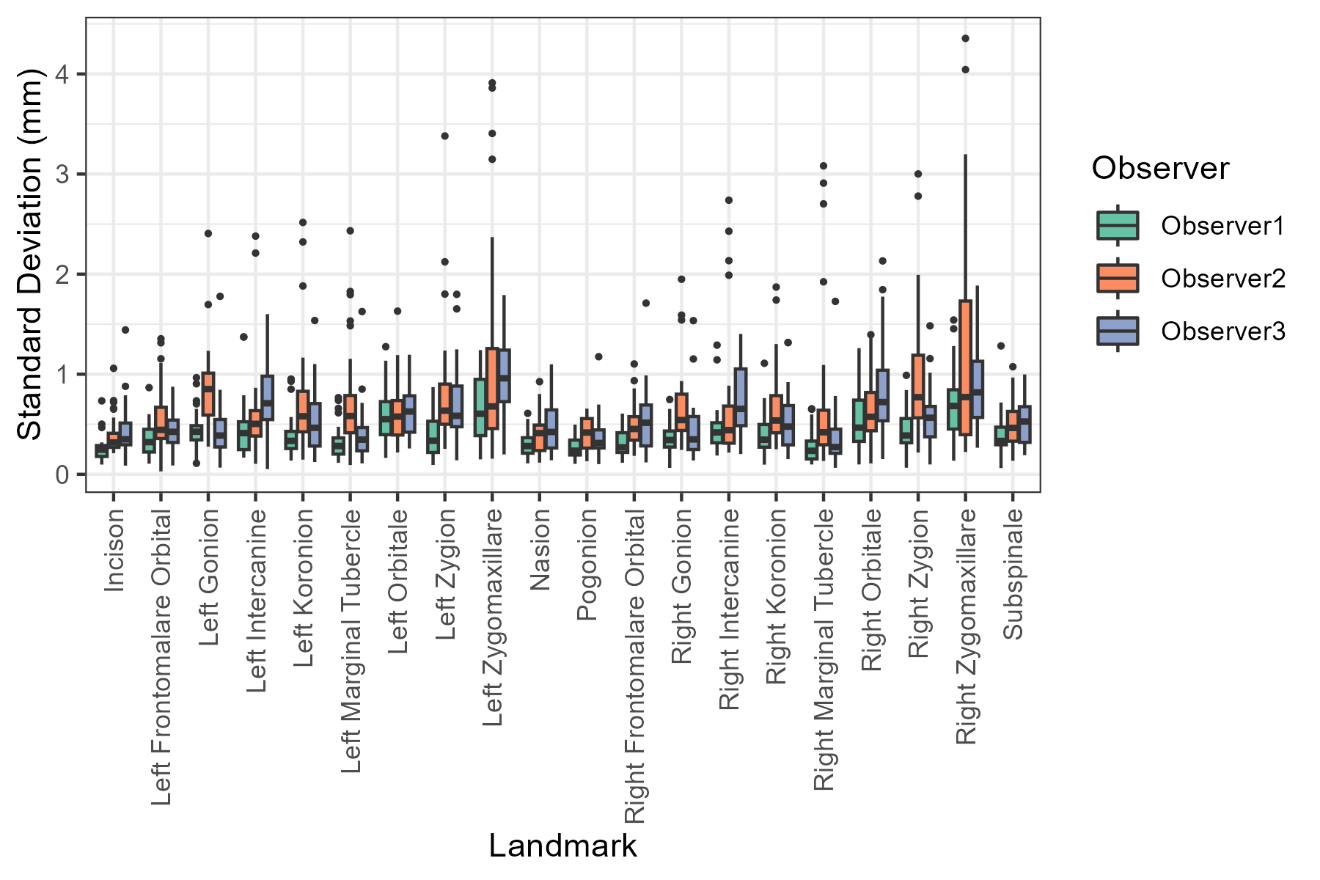
**Supplementary Figure S6:** Boxplot of intra- and inter-observer errors shows as the standard deviations in mm over the three landmarking iterations/observers averaged over the three axes per landmark.

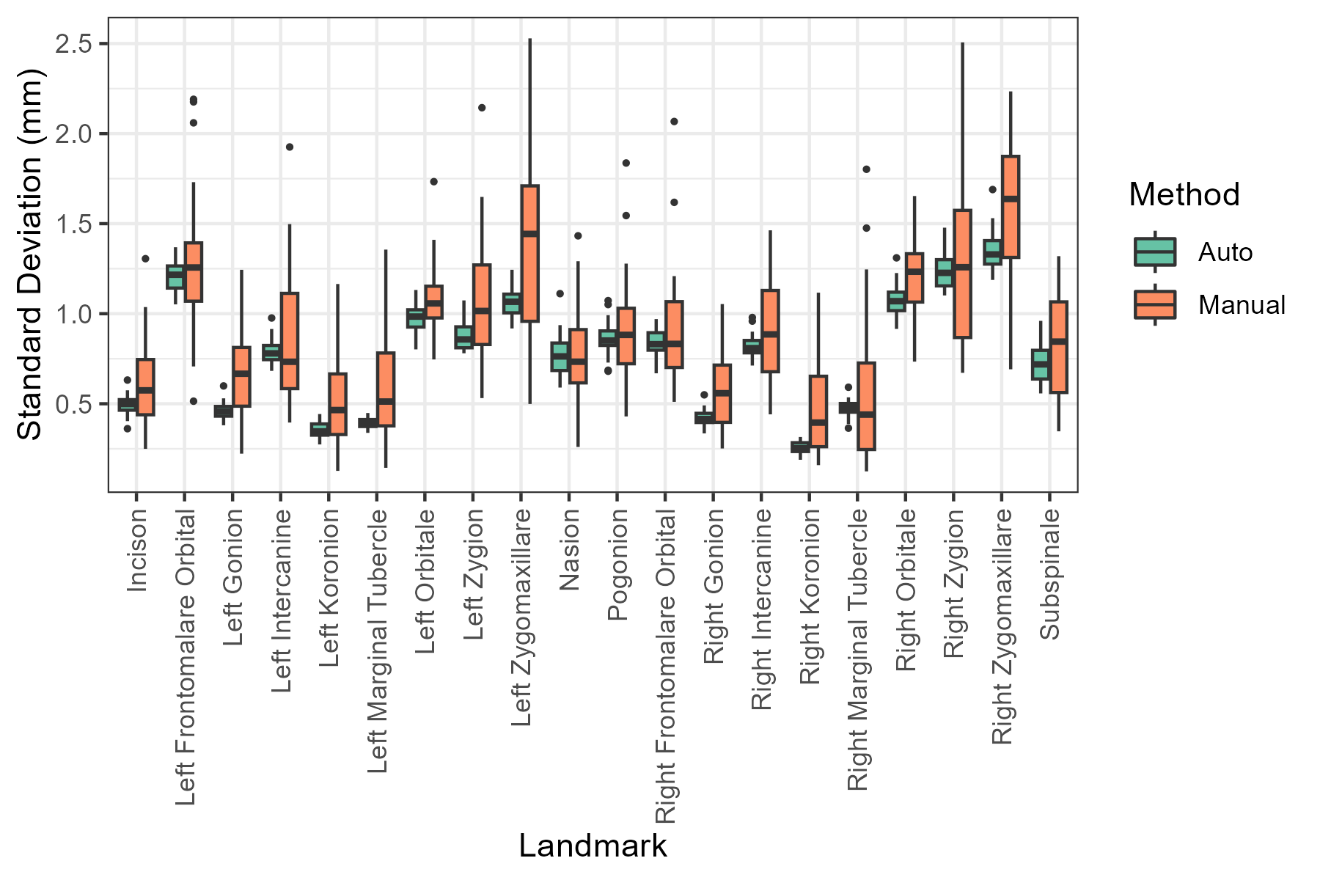

**Supplementary Table S4:** ANOVA on the centroid sizes for all manual landmarks. Skull, observer, and nested observer/iteration were inputted as factors.

|  | **Df** | **Sum Sq** | **Mean Sq** | **F value** | **Pr(>F)** |
| --- | --- | --- | --- | --- | --- |
| **Skull** | 30 | 34417.448 | 1147.248 | 1507.151 | <0.001 |
| **Observer** | 2 | 379.337 | 189.669 | 249.170 | <0.001 |
| **Skull*Observer** | 60 | 105.202 | 1.753 | 2.303 | <0.001 |
| **Observer*Iteration** | 6 | 23.672 | 3.945 | 5.183 | <0.001 |
| **Residuals** | 180 | 137.017 | 0.761 |  |  |

**Supplementary Table S5:** MANOVA on the GPA aligned manual and auto landmarks. Skull, observer, and nested observer/iteration (for manual only) were inputted as predictors.

| **MANUAL** | **Df** | **SS** | **MS** | **Rsq** | **F** | **Z** | **Pr(>F)** |
| --- | --- | --- | --- | --- | --- | --- | --- |
| **Observer** | 2 | 0.0000 | 0.0000 | 0.0000 | 0.000 | -9.801 | 1 |
| **Skull** | 30 | 0.7195 | 0.0240 | 0.9396 | 96.396 | 5.409 | 0.01 |
| **Observer*Skull** | 60 | 0.0000 | 0.0000 | 0.0000 | 0.000 | -9.950 | 1 |
| **Residuals** | 186 | 0.0463 | 0.0002 | 0.0604 |  |  |  |
| **Total** | 278 | 0.7658 |  |  |  |  |  |

| **AUTOMATIC** | **Df** | **SS** | **MS** | **Rsq** | **F** | **Z** | **Pr(>F)** |
| --- | --- | --- | --- | --- | --- | --- | --- |
| **Observer** | 2 | 0.0448 | 0.0224 | 0.1946 | 1254.732 | 7.908 | 0.01 |
| **Skull** | 30 | 0.1841 | 0.0061 | 0.8007 | 344.108 | 9.915 | 0.01 |
| **Residuals** | 60 | 0.0011 | 0.0000 | 0.0047 |  |  |  |
| **Total** | 92 | 0.2300 |  |  |  |  |  |

**Supplementary Table S6:** Inter-observer error shown as the standard deviation of the landmarks along the x, y, and z axes per landmark. Values are the standard deviation in mm. Levene’s Test shows the difference in error variance using the mean landmarking standard deviation over all axes.

|  | **Automatic** | | | | **Manual** | | | | **Levene Test** | |
| --- | --- | --- | --- | --- | --- | --- | --- | --- | --- | --- |
| **Landmark** | **X** | **Y** | **Z** | **Mean** | **X** | **Y** | **Z** | **Mean** | **F Value** | **P Value** |
| **Nasion** | 0.116 | 1.979 | 0.215 | 0.770 | 0.202 | 1.937 | 0.229 | 0.789 | 10.583 | 0.002 |
| **Subspinale** | 0.080 | 1.575 | 0.498 | 0.718 | 0.279 | 1.721 | 0.490 | 0.830 | 36.577 | <0.001 |
| **Incison** | 0.224 | 0.518 | 0.733 | 0.492 | 0.398 | 0.638 | 0.741 | 0.593 | 23.923 | <0.001 |
| **Pogonion** | 0.249 | 2.088 | 0.242 | 0.860 | 0.350 | 2.126 | 0.279 | 0.918 | 16.217 | <0.001 |
| **Right Frontomalare Orbital** | 1.464 | 0.238 | 0.816 | 0.839 | 1.447 | 0.466 | 0.848 | 0.920 | 12.605 | 0.001 |
| **Left Frontomalare Orbital** | 1.902 | 0.548 | 1.153 | 1.201 | 1.857 | 0.779 | 1.213 | 1.283 | 16.122 | <0.001 |
| **Right Orbitale** | 1.320 | 1.417 | 0.501 | 1.079 | 1.431 | 1.413 | 0.692 | 1.179 | 15.892 | <0.001 |
| **Left Orbitale** | 1.342 | 1.168 | 0.405 | 0.972 | 1.420 | 1.151 | 0.672 | 1.081 | 9.651 | 0.003 |
| **Right Zygomaxillare** | 1.602 | 0.976 | 1.465 | 1.348 | 1.974 | 1.065 | 1.702 | 1.580 | 28.742 | <0.001 |
| **Left Zygomaxillare** | 0.809 | 1.006 | 1.380 | 1.065 | 1.481 | 1.103 | 1.488 | 1.357 | 44.343 | <0.001 |
| **Right Intercanine** | 1.161 | 0.295 | 1.008 | 0.821 | 1.239 | 0.414 | 1.152 | 0.935 | 36.386 | <0.001 |
| **Left Intercanine** | 1.120 | 0.547 | 0.697 | 0.788 | 1.174 | 0.605 | 0.829 | 0.869 | 27.672 | <0.001 |
| **Right Marginal Tubercle** | 0.236 | 0.728 | 0.462 | 0.476 | 0.274 | 0.836 | 0.544 | 0.551 | 18.911 | <0.001 |
| **Right Zygion** | 1.195 | 1.039 | 1.451 | 1.228 | 1.243 | 1.048 | 1.567 | 1.286 | 35.860 | <0.001 |
| **Right Koronion** | 0.168 | 0.140 | 0.466 | 0.258 | 0.371 | 0.377 | 0.692 | 0.480 | 28.232 | <0.001 |
| **Right Gonion** | 0.055 | 0.571 | 0.638 | 0.422 | 0.155 | 0.770 | 0.764 | 0.563 | 44.009 | <0.001 |
| **Left Marginal Tubercle** | 0.144 | 0.449 | 0.589 | 0.394 | 0.244 | 0.835 | 0.692 | 0.590 | 27.754 | <0.001 |
| **Left Zygion** | 0.939 | 0.963 | 0.717 | 0.873 | 1.063 | 0.906 | 1.213 | 1.061 | 21.518 | <0.001 |
| **Left Koronion** | 0.186 | 0.184 | 0.697 | 0.356 | 0.349 | 0.389 | 0.841 | 0.526 | 26.834 | <0.001 |
| **Left Gonion** | 0.071 | 0.694 | 0.612 | 0.459 | 0.165 | 0.939 | 0.858 | 0.654 | 41.990 | <0.001 |
| **Mean** | 0.719 | 0.856 | 0.737 | 0.771 | 0.856 | 0.976 | 0.875 | 0.902 |  |  |

**Supplementary Figure S7:** Bland-Altman plots showing the Euclidean Difference in mm between manual and automatic landmarks for each axes.

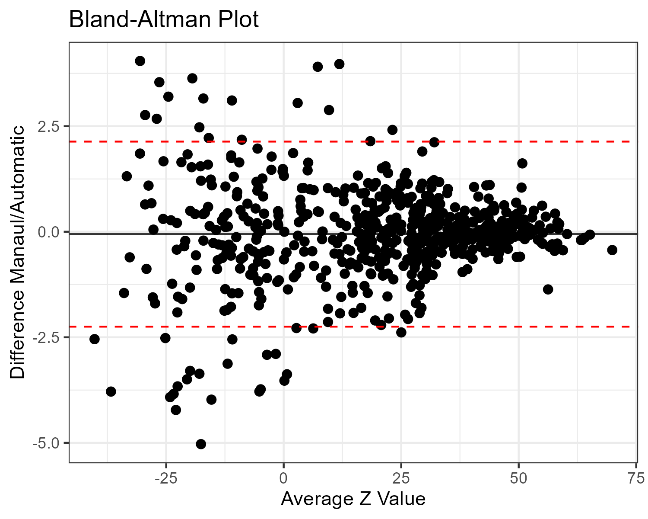

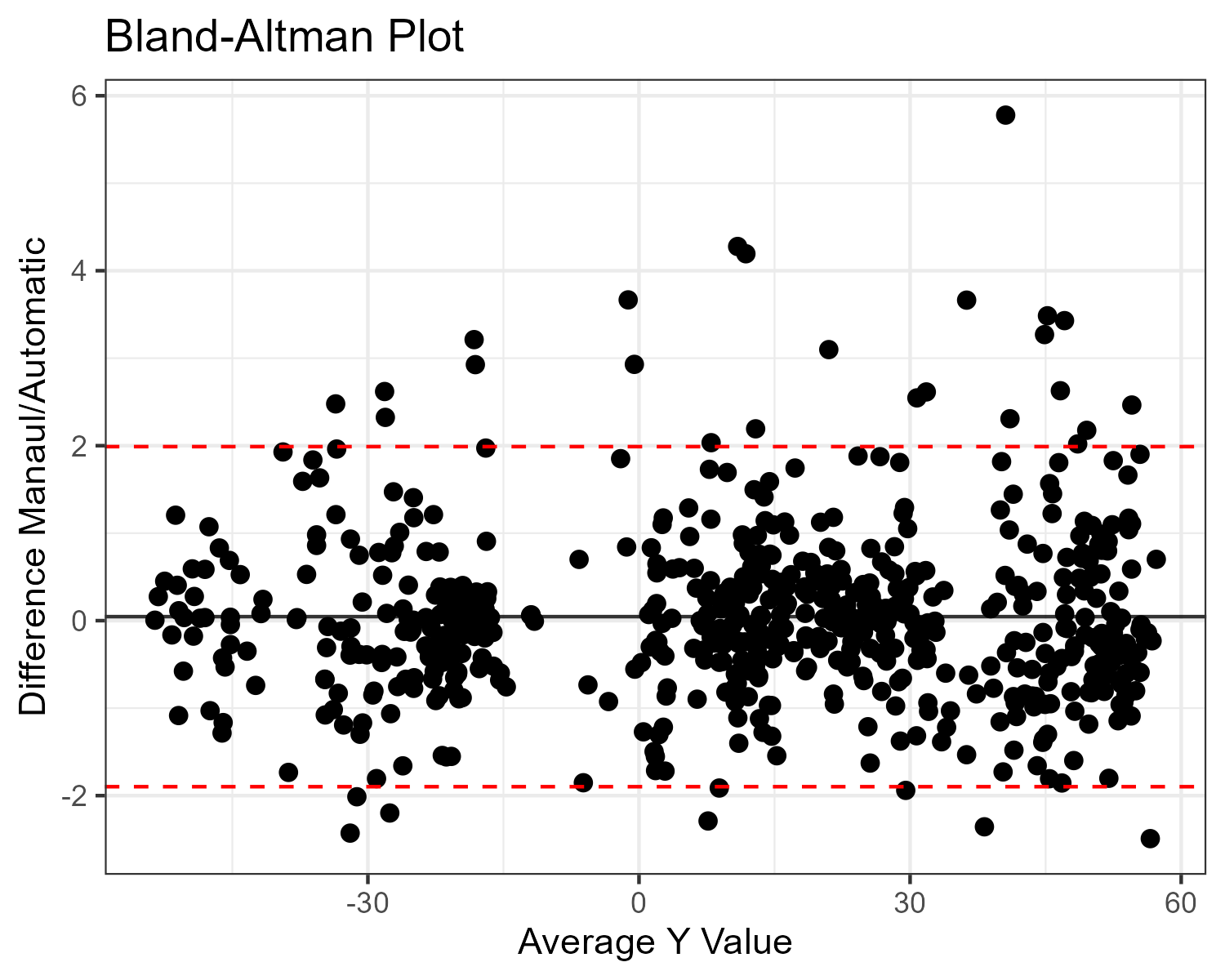

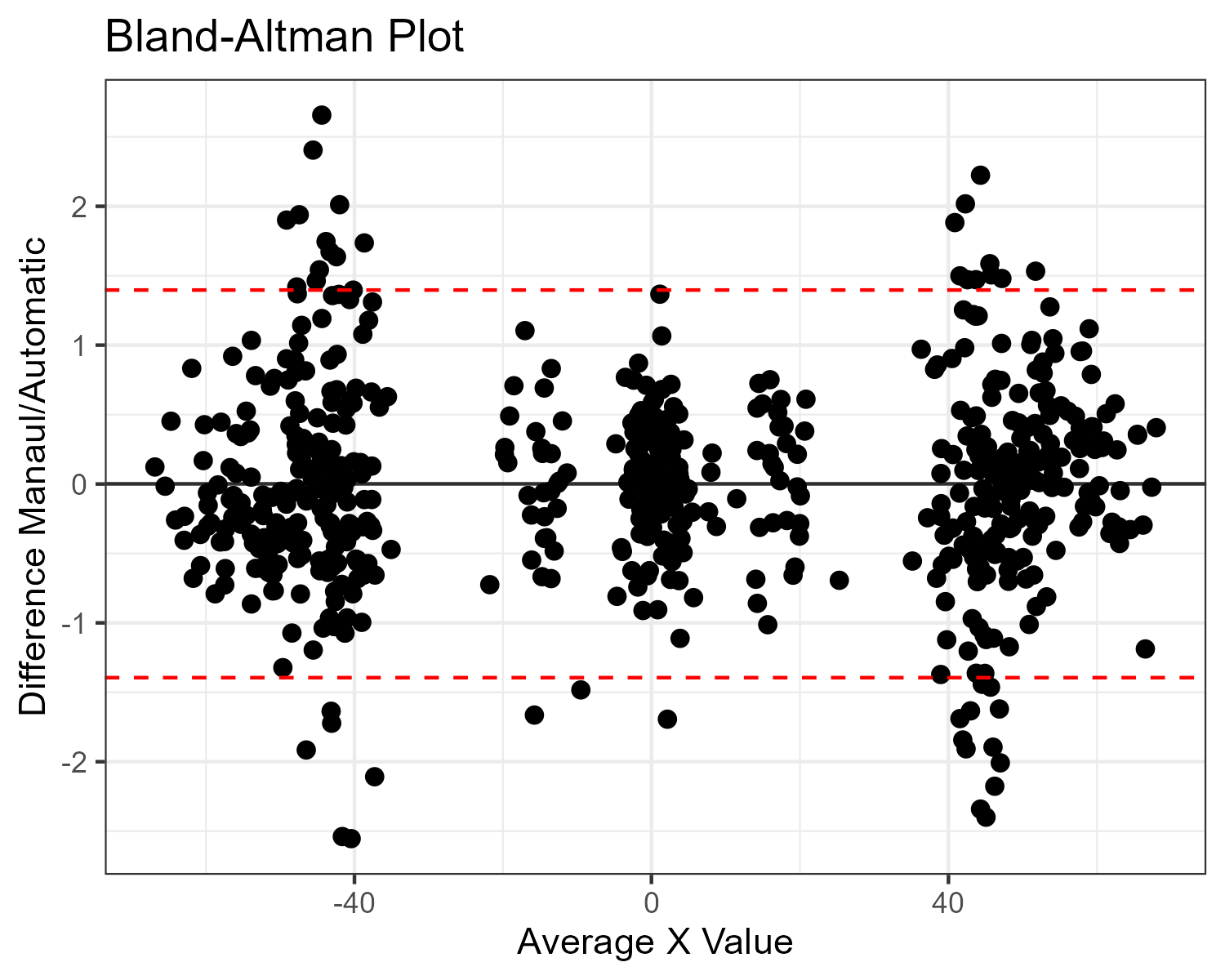

**Supplementary Figure S8:** Visual and tabular variation of the Euclidean distance error over the 20 manually placed landmarks between manual and automatic landmarks. Variation is shown as the three largest dimensions and in which direction this variation occured.

| **Landmark** | **Red** | **Blue** | **Green** |
| --- | --- | --- | --- |
| Nasion | 1.168 | 0.422 | 0.273 |
| Subspinale | 1.629 | 0.673 | 0.343 |
| Incison | 0.813 | 0.525 | 0.418 |
| Pogonion | 0.876 | 0.426 | 0.253 |
| Right Frontomalare Orbital | 1.479 | 0.560 | 0.241 |
| Left Frontomalare Orbital | 1.215 | 0.605 | 0.239 |
| Right Orbitale | 1.311 | 0.807 | 0.266 |
| Left Orbitale | 1.614 | 0.662 | 0.231 |
| Right Zygomaxillare | 2.071 | 0.719 | 0.402 |
| Left Zygomaxillare | 2.057 | 0.844 | 0.351 |
| Right Intercanine | 1.107 | 0.606 | 0.315 |
| Left Intercanine | 1.028 | 0.730 | 0.333 |
| Right Marginal Tubercle | 2.036 | 0.661 | 0.449 |
| Right Zygion | 1.533 | 0.679 | 0.266 |
| Right Koronion | 2.128 | 1.033 | 0.625 |
| Right Gonion | 1.883 | 1.227 | 0.591 |
| Left Marginal Tubercle | 2.018 | 0.610 | 0.428 |
| Left Zygion | 1.592 | 0.689 | 0.261 |
| Left Koronion | 2.101 | 1.232 | 0.595 |
| Left Gonion | 1.816 | 1.151 | 0.393 |

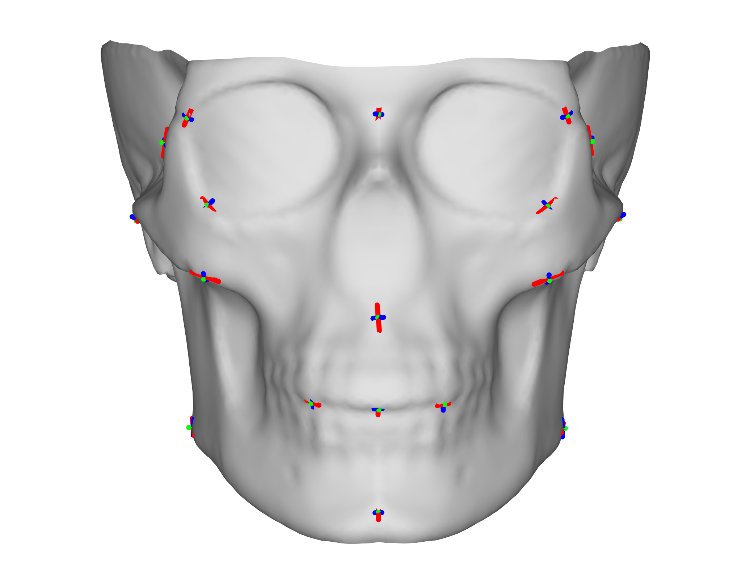

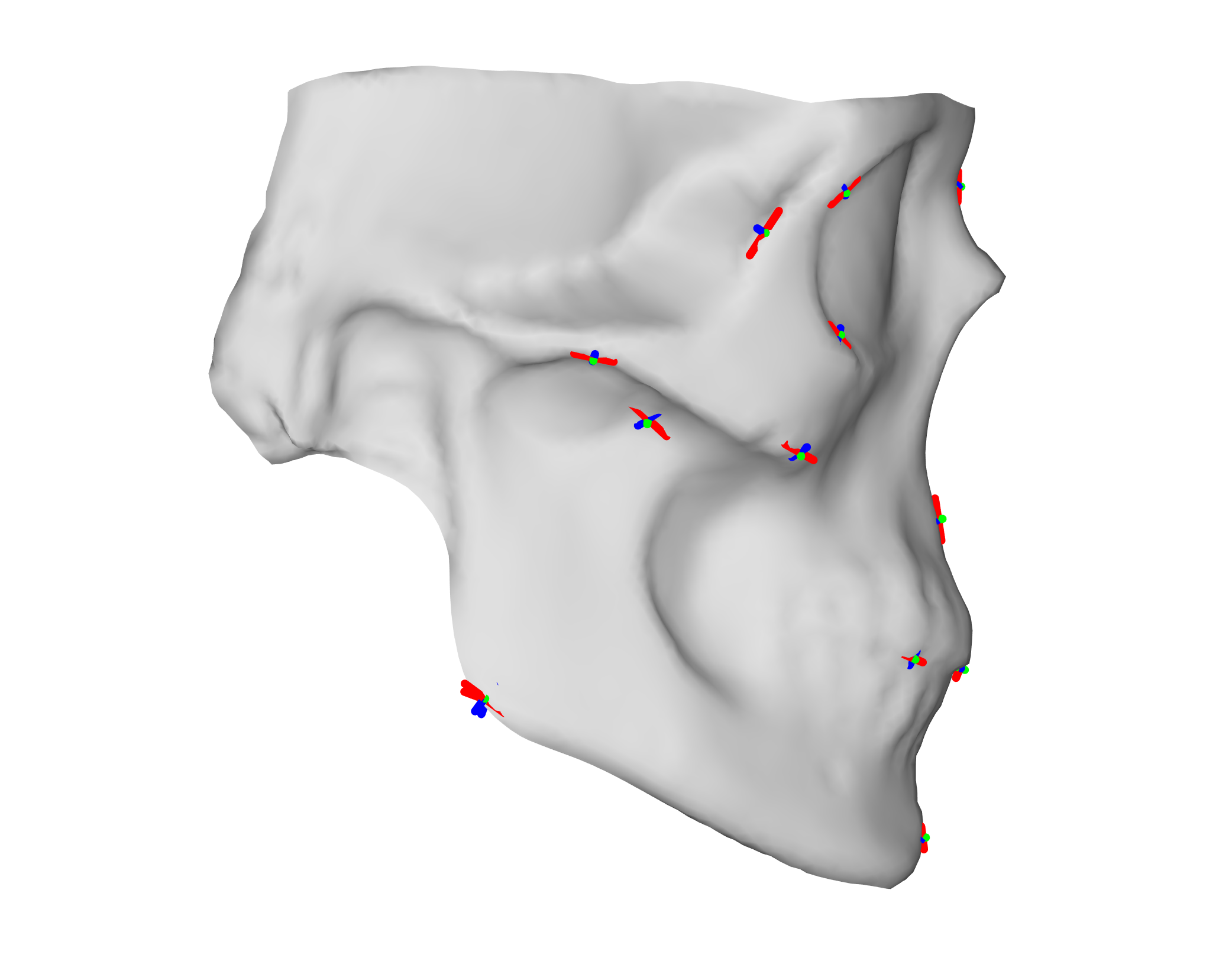

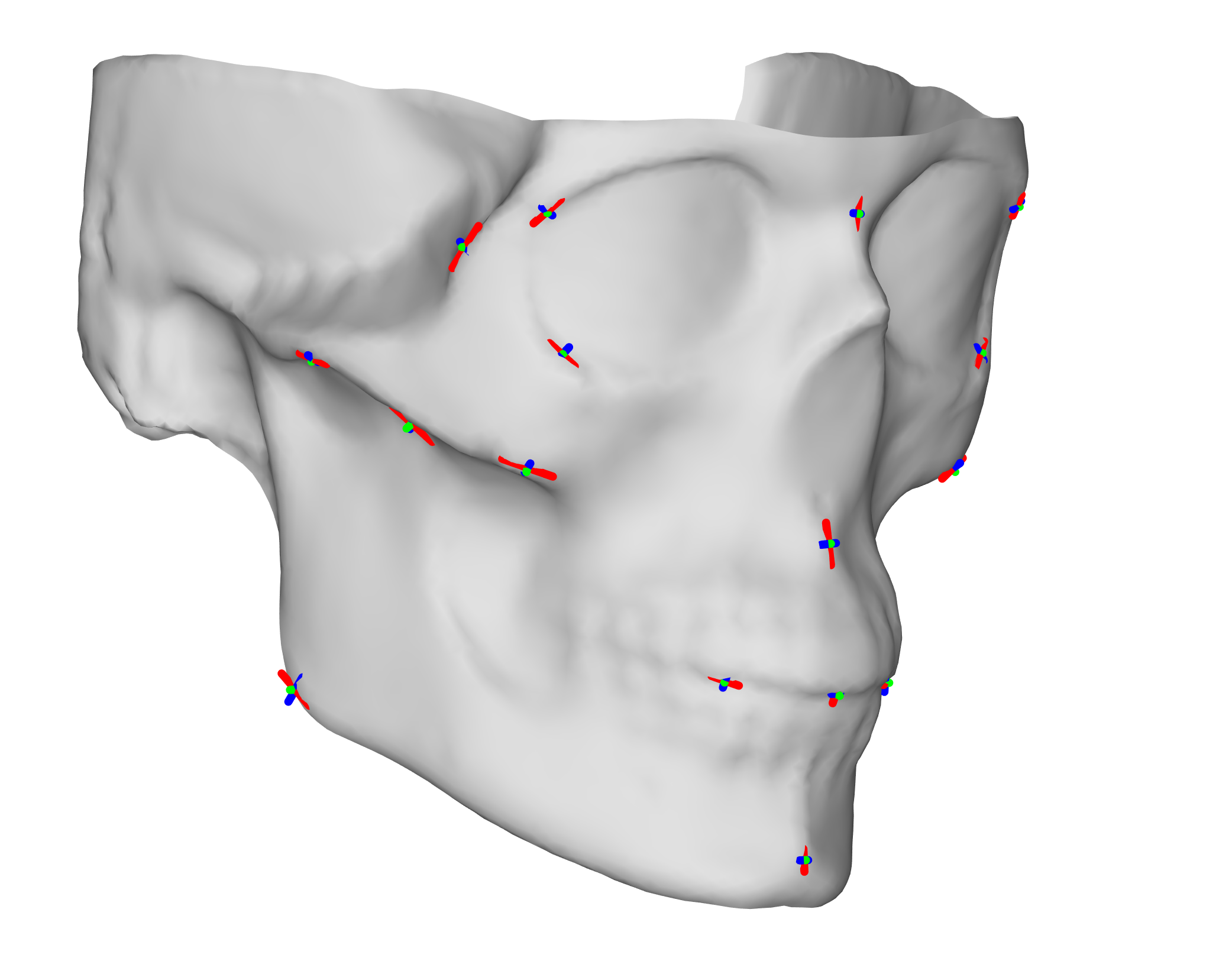

|  | **Df** | **SS** | **MS** | **Rsq** | **F** | **Z** | **Pr(>F)** |
| --- | --- | --- | --- | --- | --- | --- | --- |
| **Observer** | 2 | 0.0224 | 0.0112 | 0.0487 | 5.606 | 4.773 | 0.01 |
| **Skull** | 30 | 0.1862 | 0.0062 | 0.4049 | 3.109 | 8.751 | 0.01 |
| **Method** | 1 | 0.0242 | 0.0242 | 0.0525 | 12.102 | 5.077 | 0.01 |
| **Residuals** | 152 | 0.2272 | 0.0027 | 0.4939 |  |  |  |
| **Total** | 185 | 0.4600 |  |  |  |  |  |

**Supplementary Table S7:** MANOVA on the GPA aligned landmarks. Skull, observer, and method were inputted as predictors.

|  | **Df** | **Pillai** | **approx F** | **num Df** | **den Df** | **Pr(>F)** |
| --- | --- | --- | --- | --- | --- | --- |
| **Observer** | 1 | 0.7909 | 48.3050 | 13 | 166 | <0.001 |
| **Skull** | 1 | 0.3413 | 6.6176 | 13 | 166 | <0.001 |
| **Method** | 1 | 0.0239 | 0.3132 | 13 | 166 | 0.99 |
| **Observer*Skull** | 1 | 0.0035 | 0.0447 | 13 | 166 | 1 |
| **Observer*Method** | 1 | 0.0005 | 0.0069 | 13 | 166 | 1 |
| **Skull*Method** | 1 | 0.0262 | 0.3429 | 13 | 166 | 0.98 |
| **Residuals** | 178 |  |  |  |  |  |

**Supplementary Table S8:** MANOVA on the first 13 PCs of the auto and manual landmarks (explaining ~95% of the variation). Skull, observer, and method were inputted as predictors.

**Supplementary Figure S7:** Visualization of the masking process on a CT scan. A: cleaned CT scan from MUG500+, B: After shrinkwrapping, C: After decimation/remeshing, D: After masking.

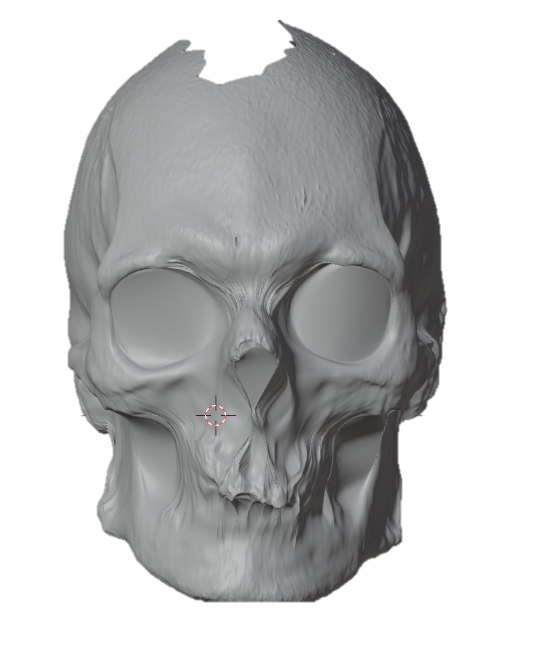

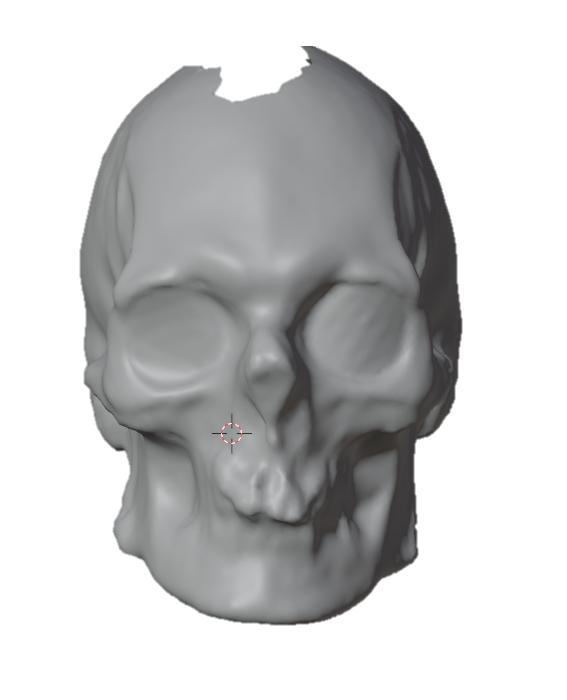

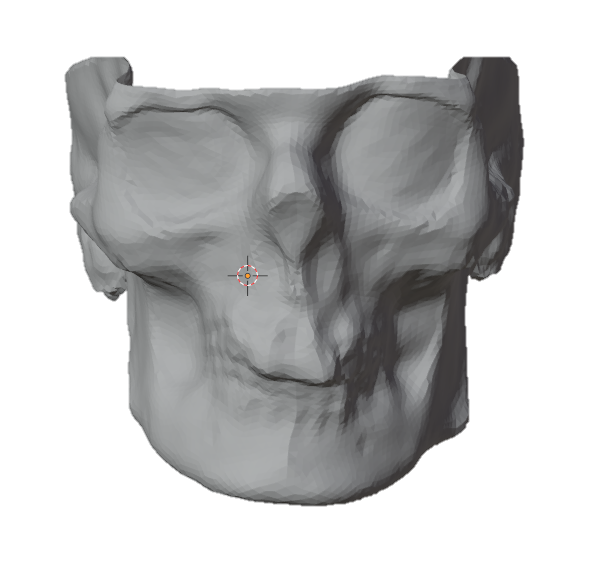

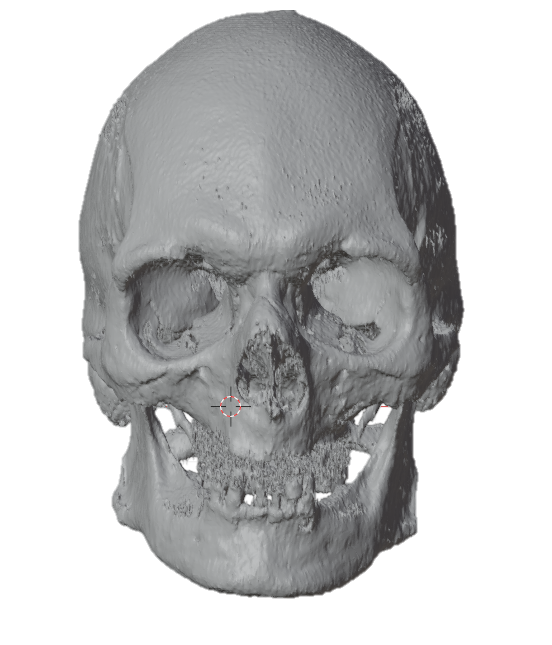

A

B

C

D
